# Molecular mechanisms underlying the extreme mechanical anisotropy of the flaviviral exoribonuclease-resistant RNAs (xrRNAs)

**DOI:** 10.1101/2020.05.26.117747

**Authors:** Xiaolin Niu, Qiuhan Liu, Zhonghe Xu, Zhifeng Chen, Linghui Xu, Lilei Xu, Jinghong Li, Xianyang Fang

## Abstract

Mechanical anisotropy is an essential property for many biomolecules to assume their structures, functions and applications, however, the mechanisms for their direction-dependent mechanical responses remain elusive. Herein, by using single-molecule nanopore sensing technique, we explore the mechanisms of directional mechanical stability of the xrRNA1 RNA from ZIKA virus (ZIKV), which forms a complex ring-like architecture. We reveal extreme mechanical anisotropy in ZIKV xrRNA1 which highly depends on Mg^2+^ and the key tertiary interactions. The absence of Mg^2+^ and disruption of the key tertiary interactions strongly affect the structural integrity and attenuate mechanical anisotropy. The significance of ring structure in RNA mechanical anisotropy is further supported by steered molecular dynamics simulations on ZIKV xrRNA1 and another two RNAs with ring structures, the HCV IRES and THF riboswitch. We anticipate the ring structures can be used as key elements to build RNA-based nanostructures with controllable mechanical anisotropy for biomaterial and biomedical applications.

## INTRODUCTION

RNAs’ diverse roles in many cellular processes are dictated by their propensities to fold into stable three-dimensional structures driven by numerous tertiary interactions^1, 2^. These tertiary interactions are important to RNAs’ structure, stability, dynamics, and folding kinetics^3^. While RNA folds into stable structure as it is synthesized, it undergoes unfolding and refolding events during many cellular processes such as translation, replication, reverse transcription, etc., during which molecular motors exert forces on the RNA in a directional manner (i.e. ribosome proceeds along the RNA template in the 5’→3’ direction)^4^. For many nucleic acids, mechanical anisotropy is an inherent property which mechanical behavior varies with the direction of applied forces and is of high biological significance. For instance, the three-wayjunction-pRNA derived from φ29 DNA packaging motor has been shown to exhibit high mechanical anisotropy upon Mg^2+^ binding, which relates to its capability to withstand the strain caused by DNA condensation^5, 6^. The mechanical anisotropy of the human telomeric DNA G-quadruplex can be changed by ligand binding, corresponding to its regulation on replication or transcription^7^. Understanding how RNA responds to a mechanical stretching force and the molecular mechanisms defining its mechanical anisotropy is important for not only elucidating key principles governing various mechano-biological processes but developing novel RNA-based biomaterials with tailored mechanical properties and RNA-targeted therapeutics^8^, thus, has been an important research topic in the field of RNA mechanics^9^.

Exoribonuclease-resistant RNAs (xrRNAs) are a group of RNA elements which are capable of resisting the degradation by exonuclease^10, 11^. The ability to resist Xrn1 is surprising as Xrn1 is capable of processively degrading highly structured RNAs^12^. Recent crystal structures of xrRNAs from Murray Valley encephalitis virus (MVEV, xrRNA2)^13^ and ZIKV (xrRNA1)^14^ reveal a stable and compact RNA fold centered on a three-way junction that forms an unusual ringlike structure through which the 5’ end passes (**Figure 1a and b**). These structures suggest a “molecular brace” model for high resistance to directional degradation by the 5’-3’ exonucleases including Xrn1, Dxo1 and RNase J1, etc^15, 16^. Upon encountering the xrRNAs, Xrn1 must pull the 5’ end of the RNA through the ring, therefore creating a mechanical unfolding problem that the enzyme can’t resolve. In contrast, the viral RNA-dependent RNA polymerase (RdRP) approaching in the 3’→5’ direction during (-) strand synthesis can readily traverse this structure (**Figure 1c**). It is hypothesized that the xrRNAs exhibit significant mechanical anisotropy^13, 16^, which however has not been experimentally proved and the underlying mechanisms remain largely unknown.

**Figure 1.**
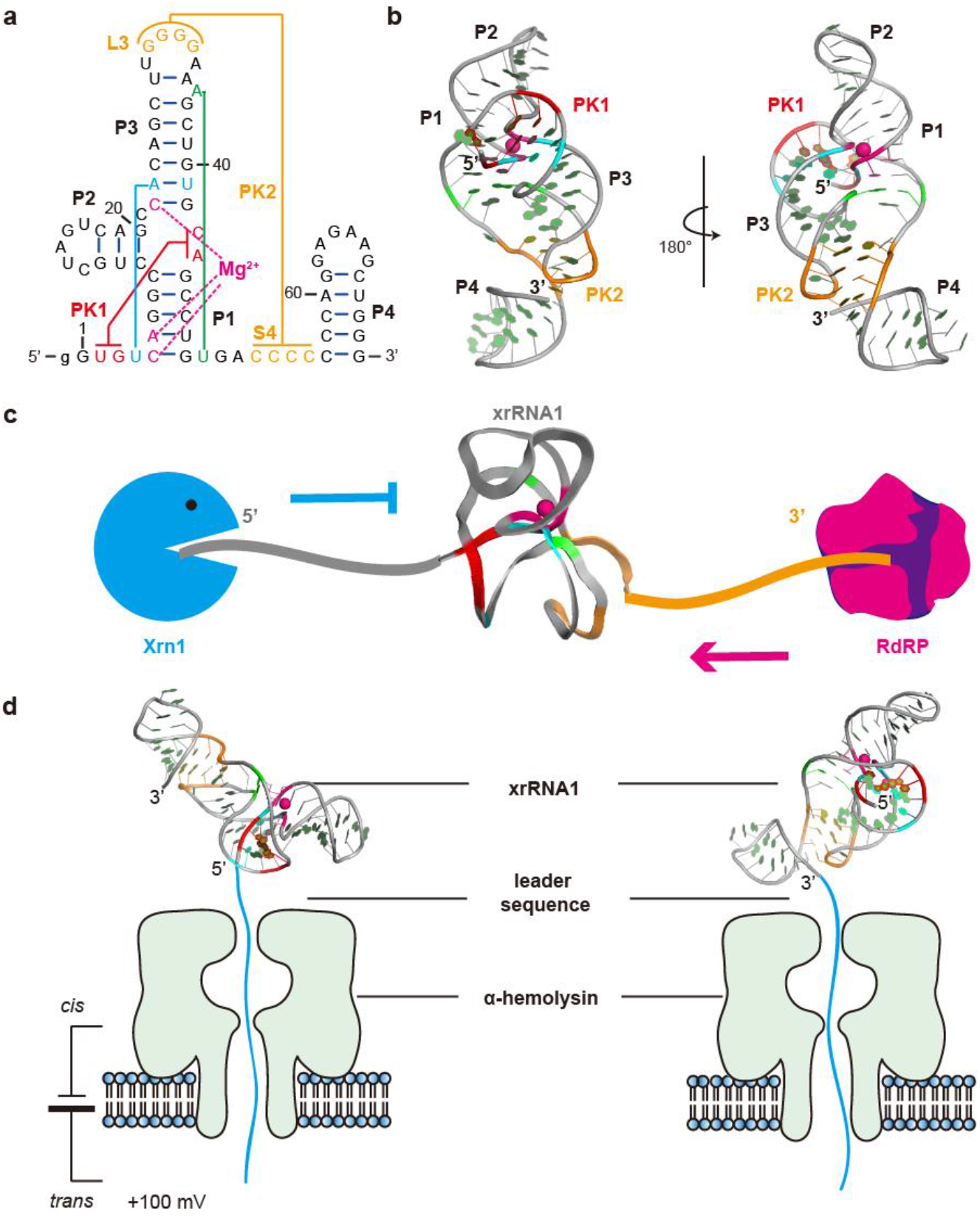
Mimicking directional unfolding of ZIKV xrRNA1 in the α-HL nanopore. (**a, b**) Secondary (**a**) and tertiary structure (**b**) of ZIKV xrRNA1 (PDB code: 5TPY), which contains several tertiary interactions (cyan: U4:A23:U41, green: A36:U50) and two pseudoknots, PK1 (red, U2G3:C43G44) and PK2 (orange, G30-G33:C53-C56). Additionally, one Mg^2+^ is found to coordinate with the phosphate of C5, A6 and C22 (pink). (**c**) A cartoon shows that xrRNA1 can resist the degradation by exoribonuclease Xrn1 from 5’→3’ direction, yet be traversed by the viral RdRP from 3’→5’ direction. (**d**) Cartoons showing the directional unfolding and translocation of ZIKV xrRNA1 through the α-HL nanopore, where a 36-nt poly(rA) leader sequence was co-transcribed with xrRNA1 at the 5’- or 3’-end to facilitate its translocation through the nanopore.

Various single-molecule force spectroscopy techniques, e.g., optical tweezers, magnetic tweezers and atomic force microscope (AFM) have been successfully applied to study RNA mechanical unfolding^7–8, 17^. By pulling the RNAs from both ends, these approaches can identify the intermediate states along the folding pathway, but they also have obvious limitations in which the unfolding process is not unidirectional, differ from that *in vivo*. Fortunately, recent developments in single-molecule nanopore sensing technology have allowed the unfolding and translocation of RNAs through biological pores in a defined direction to be observed with fine details^18, 19, 20^. By taking advantage of the ability to electrically detect charged biomolecules through a nanometer-wide channel, several types of nanopores including α-hemolysin (α-HL) nanopore have been developed^21, 22^. The narrowest constriction of α-hemolysin is 1.4 nm in the stem domain (**Figure S1**), and thus, it only allows the translocation of single-stranded nucleic acids, but folded nucleic acids must unfold to pass through the pore^21^.

In this work, using the α-hemolysin nanopore sensing technique, we investigate the directional mechanical stability of the xrRNA1 from ZIKV at single-molecule level. A 36-nt poly(rA) leader sequence is co-transcribed at the 5’ or 3’ end of ZIKV xrRNA1, respectively, to direct the translocation through the nanopore (**Figure 1d**). The duration time that xrRNA1 is trapped in the nanocavity before unraveling provides a measure of the structure’s mechanical stability. Extreme mechanical anisotropy is observed for ZIKV xrRNA1 which highly depends on the presence of Mg^2+^ and the tertiary interactions that stabilize the ring-like topological structure. The absence of Mg^2+^ and disruption of the key tertiary interactions strongly affect the structural integrity of the ring structure and attenuate mechanical anisotropy. The correlation is further supported by steered molecular dynamics simulation (SMD) analysis on ZIKV xrRNA1 and another two RNAs with ring-like architecture, including the core domain of the Hepatitis C virus internal ribosome entry site^23^ (HCV IRES) and the holo-aptamer domain of the tetrahydrofolate (THF) riboswitch^24^, suggesting an important role of the ring-like architectures in defining the respective RNAs’ mechanical anisotropy.

## RESULTS

### ZIKV xrRNA1 exhibits extreme mechanical anisotropy

Mg^2+^ ions are known to be important for the structure and stability of RNA molecules^25^. As shown in **Figure 1a-b**, a Mg^2+^ ion is found to coordinate with the phosphates of C5, A6 and C22 in the crystal structure of ZIKV xrRNA1, implying that Mg^2+^ ions may play an important role in the folding of xrRNA1. To better understand how Mg^2+^ affect the overall structure and stability of ZIKV xrRNA1, small angle x-ray scattering (SAXS) and differential scanning calorimetry (DSC) experiments were performed (further details are provided in the Supplementary Information). SAXS data indicates a Mg^2+^-induced structural transition between the unfolded and folded states, and the midpoint of the transition is about 0.3 mM Mg^2+^ (**Figure S2**). DSC experiments demonstrate the cooperative unfolding of xrRNA1 highly depends on Mg^2+^ concentrations (**Figure S3**). Both experiments show that xrRNA1 is fully folded in 5 mM Mg^2+^, and the high salt concentration (1M KCl) has a minor effect on the structure of xrRNA1. The following single-molecule nanopore sensing experiments are firstly conducted in 1M KCl electrolyte solutions containing 5 mM Mg^2+^.

To facilitate directional translocation of ZIKV xrRNA1 through the α-hemolysin (α-HL) nanopore, a 36-nt poly(rA) leader sequence is co-transcribed with the xrRNA1 at the 5’ or 3’ end (hereafter as 5’_A36_-xrRNA1 and 3’_A36_-xrRNA1, respectively) (**Figure 2a and b**). Similar designs have been used to facilitate directional translocation of the human telomere i-motif DNA and T2 pseudoknot RNA through α-HL nanopore^19, 26^. DSC experiments on these xrRNA1s show that the leader sequence has minor effects on the thermal stability of xrRNA1s (**Figure S4, Table S2)**. Under a transmembrane voltage, the single-stranded RNA leader sequence (36nt, ~ 18 nm) which is much longer than the nanopore passage (10 nm), will be captured and occupy the entire nanopore channel first, then guides the directional unfolding and translocation of the downstream structured RNA elements through the nanopore and releases into the trans solution(**Figure S5**). While being captured and translocated through the confined space of the nanopore, the RNAs are subjected to significant electric force and undergo structural changes then unfolding, the ions occupying the nanopore are excluded, therefore results in various characteristic current blockade signatures characterized by amplitude and duration time.

**Figure 2.**
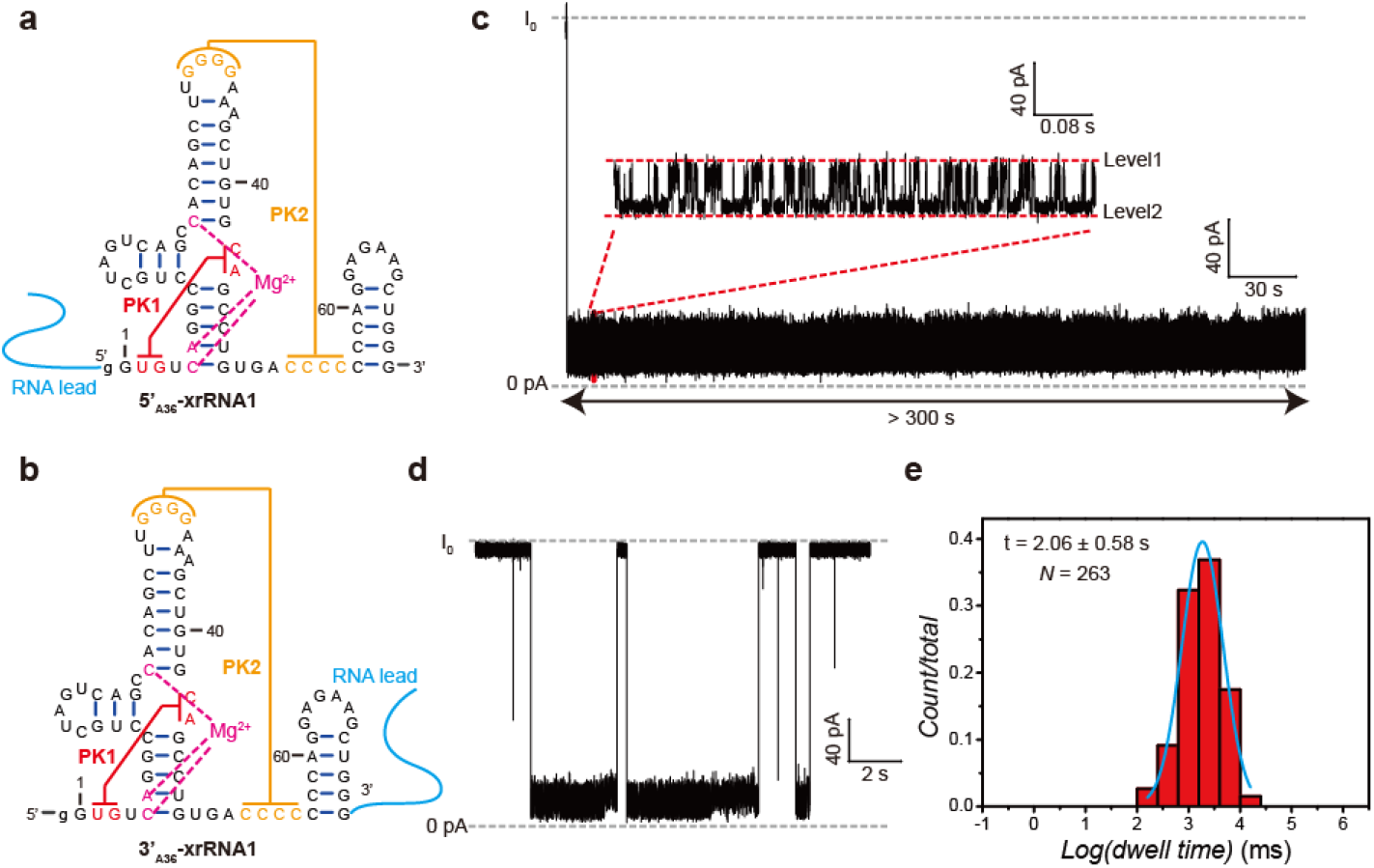
ZIKV xrRNA1 exhibits significant mechanical anisotropy in the presence of 5 mM Mg^2+^. (**a, b**) Secondary structure of 5’_A36_-xrRNA1 (**a**) and 3’_A36_-xrRNA1 (**b**), which a 36-nt poly(A) was co-transcribed at the 5’- or 3’- end of the xrRNA1. (**c, d**) Representative current blockade traces for unfolding of 5’_A36_-xrRNA1 (**c**) and 3’_A36_-xrRNA1 (**d**) in the α-hemolysin nanopore. (**e**) Representative dwell time distribution histogram for the unfolding of 3’_A36_-xrRNA1. The dwell time were the mean value calculated from N > 200 events and 3 independent repeat experiments. All the experiments were conducted at +100 mV in a buffer containing 20 mM Tris, 1 M KCl and 5 mM MgCl_2_ at 25℃.

Initial nanopore sensing studies were conducted for the ZIKV xrRNA1 sequences in 1M KCl supplemented with 5 mM MgCl_2_ (cis side) at +100 mV. Under such condition, the force applied to leader sequence is around 8 pN^19^, close to that exerted by molecular machines^27, 28^. **Figure 2c** shows a representative current blockade signature for the 5’_A36_-xrRNA1 through the nanopore, which only very long current blockade is observed. The initial drop of the ionic current to less than 20% of the open-pore current (I_0_) is interpreted as entry of the 5’ RNA leader sequence into the pore and occupation of the β–barrel channel, then the current switches frequently between two different levels with the relative conductance (I/I_0_) of 13% (Level 1) and 6% (Level 2), respectively. This may be due to the different orientations of xrRNA1 when threaded within the vestibule of the pore, and similar phenomena were observed in the previous research^29^. After several attempts, we can’t observe any complete 5’_A36_-xrRNA1 unfolding and translocation event during 300 s time window, which is limited by vulnerability of biological nanopore system^30^. Further increasing the transmembrane voltage to 160 or 180 mV shows similar current blockade signatures (**Figure S6**). These initial observations suggest that the kinetics of 5’_A36_-xrRNA1 unfolding and translocation through the α-HL nanopore is too slow to be monitored in a reasonable time window. In contrast, 3’_A36_-xrRNA1 in the same condition (5 mM Mg^2+^) is readily translocated through the α-HL nanopore, generating current blockade signatures distinct from that for 5’_A36_-xrRNA1 (**Figure 2d**). Analysis of >200 of the events reveals four types of current blockade signatures (**Figure S7**). Type 1 represents the most typical events and constitutes more than 85% of all the events. Moreover, the blockade signal is different from that for xrRNA1 without a leader sequence, which generates a two-stage blockade current pattern (**Figure S8**). Therefore, type1 signal is attributed to 3’→5’ directional translocation guided by the leader sequence and the following dwell time statistical analysis were all based on type1. The mean duration time for the 3’→5’ directional translocation of 3’_A36_-xrRNA1 in the nanopore is estimated as 2.06 ± 0.58 s (**Figure 2e**), much shorter than that of 5’→3’ directional translocation of 5’_A36_-xrRNA1 in the nanopore, which is longer than 300 s. Due to the complexity of xrRNA1 structure and unfolding pathway^31^, it’s difficult to have a thorough molecular interpretation of the blockade current traces, however, the duration time that xrRNA1 is trapped in the nanocavity before unraveling provides a measure for the structure’s mechanical stability^19, 29, 32^. The dramatic variation of the mean duration time of xrRNA1 between the 5’→3’ and 3’→5’ directional translocation in α-HL nanopore suggests that xrRNA1 exhibits extreme mechanical anisotropy.

### The mechanical anisotropy of ZIKV xrRNA1 highly depends on Mg^2+^

The prolonged duration time (> 300 s) for 5’_A36_-xrRNA1 in 5 mM Mg^2+^ prevents obtaining required populations of current blockages for statistical analysis, thus, mutants of xrRNA1 were designed and screened to reduce its 5’→3’ mechanical stability mildly but maintain overall structure, which results in a quadruple mutant (U2A+U28A+A36C+C53G, hereafter as xrRNA1-X) that resembles ZIKV xrRNA1 (**Figure S9**). In the crystal structure of ZIKV-xrRNA1, U2 (U2-A44) and C53 (C53-G33) are involved in long-range base-pairing interactions in the first (PK1) and second (PK2) pseudoknots formation, respectively; A36 and U50 form a reverse Watson-Crick long-range base pair that closes the ring structure to “lasso” the RNA that passes through; U28 and A35 form a Hoogsteen base pair which is adjacent to PK2 (**Figure 1a-b**). DSC experiments show that the quadruple mutation cause marginal reduction in the thermal stability of ZIKV xrRNA1, the melting temperature for xrRNA1-X is only slightly lower than ZIKV xrRNA1 (from 75.2 to 78.52 ℃) (**Figure S9c**). SAXS analysis shows that xrRNA1-X shares similar PDDF (*R_g_, D_max_*) as xrRNA1 and the experimental scattering curve of xrRNA1-X in 5 mM Mg^2+^ can fit nicely with ZIKV xrRNA1 crystal structure (**Figure S9d, 10a**). These data clearly show that the quadruple mutations in xrRNA1-X marginally reduce xrRNA1’s overall stability and cause minimal effects on its overall structure.

Both 5’_A36_-xrRNA1-X and 3’_A36_-xrRNA1-X were analyzed by nanopore sensing studies. As expected, 5’_A36-_xrRNA1-X in 5 mM Mg^2+^ readily undergoes 5’→3’ directional translocation through the α-HL nanopore at +100 mV within a measurable time window. 5’_A36_-xrRNA1-X shares some similarity in blockade current signatures as 5’_A36_-xrRNA1, for example, the leader sequence of 5’_A36_-xrRNA1-X is firstly captured by the nanopore and occupies the β-barrel channel, therefore causes rapid drop of the current to less than 20% of the open channel current, and the current switches frequently between two different levels with the relative conductance (I/I_0_) of 13% (Level 1) and 6% (Level 2), respectively (**Figure 3a**). The mean duration time of 5’_A36_-xrRNA1-X through the nanopore is estimated to be 60.78 ± 5.32 s (**Figure 3c**). The blockade current signature for 3’_A36_-xrRNA1-X is similar to that in 5 mM Mg^2+^ (**Figure 3b**), the mean duration time is estimated to be 2.00 ± 0.15 s, which is about 30 times shorter than that for 5’_A36_-xrRNA1-X (**Figure 3c**). Obviously, similar as xrRNA1, xrRNA1-X exhibits significant mechanical anisotropy in 5 mM Mg^2+^, thus, xrRNA1-X could be used to represent xrRNA1 in the subsequent quantitative analysis.

**Figure 3.**
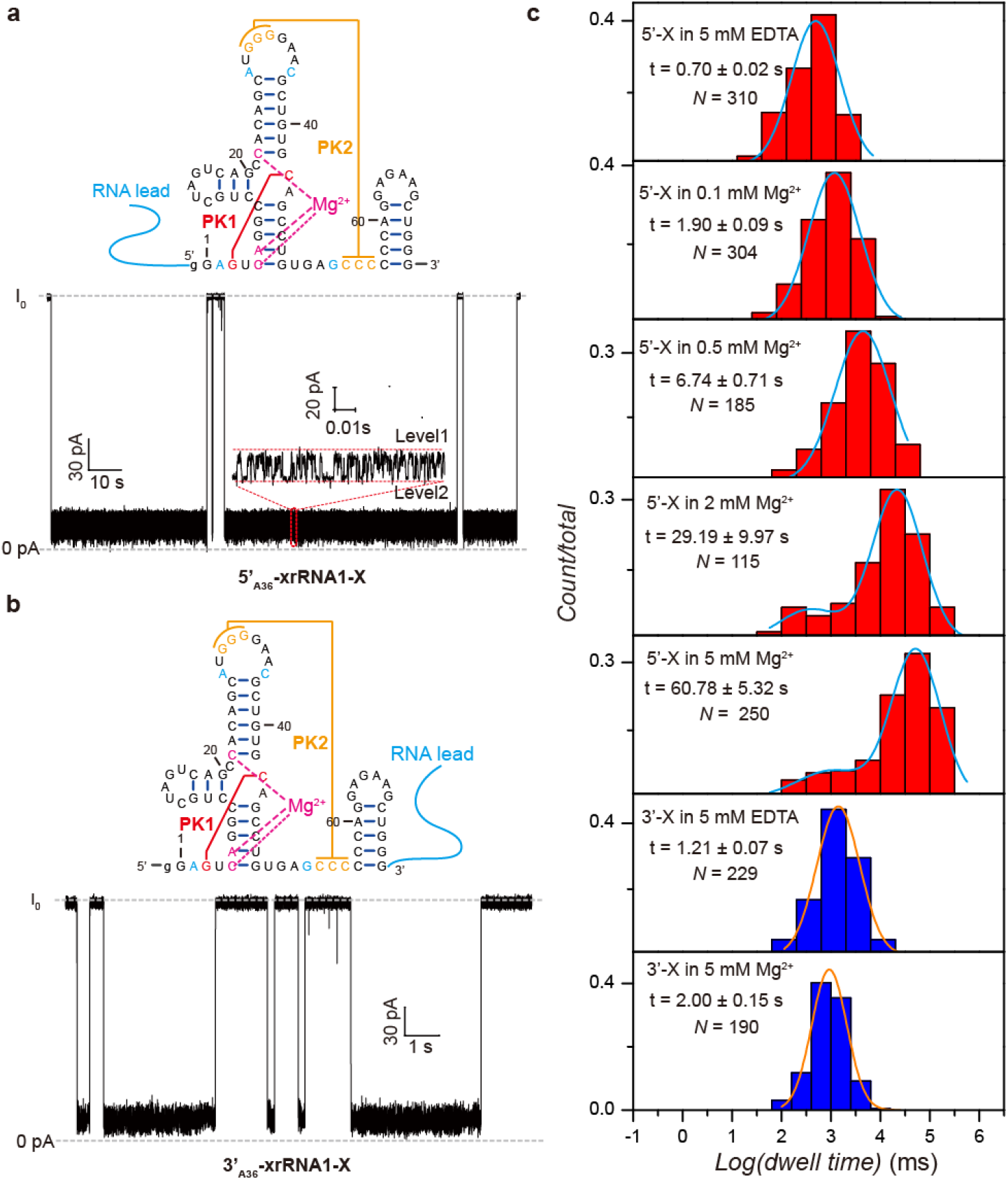
Mechanical anisotropy of ZIKV xrRNA1-X highly depends on Mg^2+^. (**a, b**) Secondary structures and representative current traces for the unfolding and translocation of 5’_A36_-xrRNA1-X (**a**) and 3’_A36_-xrRNA1-X (**b**) through the nanopore in the presence of 5 mM Mg^2+^. (**c**) Dwell time distribution histograms for 5’_A36_-xrRNA1-X (red) and 3’_A36_-xrRNA1-X (blue) in different Mg^2+^ concentrations.

It’s interesting to know how Mg^2+^ affects the directional mechanical stability of xrRNA1. To conduct the Mg^2+^-dependent directional translocation studies, 5’_A36_-xrRNA1-X was placed in electrolyte solution containing 1M KCl supplemented with various concentrations of Mg^2+^ (*cis* side). The blockade current signature for 5’_A36_-xrRNA1-X in different Mg^2+^ concentrations are very similar but the mean duration time becomes longer as Mg^2+^ concentration increases. xrRNA1 shows a well-defined distribution centered at ~0.7 s in 5 mM EDTA, as Mg^2+^ increases a second peak emerged and shifts to longer dwell times above 0.5 mM Mg^2+^, which is consistent with Mg^2+^ induced folding of xrRNA1 derived from SAXS and DSC data (**Figure 3c**). These data indicated that the 5’→3’ directional mechanical stability of xrRNA1 highly depends on Mg^2+^. In contrast, the 3’→5’ directional mechanical stability of xrRNA1 is less dependent on Mg^2+^. The mean duration time for 3’_A36_-xrRNA1-X in 5 mM EDTA is estimated to be 1.21 ± 0.1 s, only slightly smaller than that in 5 mM Mg^2+^ (2.00 ± 0.15 s) and the blockade current signatures are very similar (**Figure 3c**). Overall, the mechanical anisotropy of ZIKV xrRNA1, which is the difference between the 5’→3’ and 3’→5’ directional mechanical stability, highly depends on Mg^2+^.

### Loss of key tertiary interactions attenuates ZIKV xrRNA1 mechanical anisotropy

As shown in **Figure 1a-b**, ZIKV xrRNA1 contains a ring-like architecture centered on a three-way junction which is further stabilized by two pseudoknots^14^. The helical element in PK2 can form a continuous helix with the P4 duplex through coaxial stacking, which may further stabilize the overall fold^14^. To understand how these tertiary interaction affect the mechanical anisotropy, mutants of ZIKV xrRNA1 were generated and analyzed by nanopore sensing.

Previous studies on MVEV xrRNA2 suggested the importance of S1-S3 pseudoknot in stabilizing the active conformation of xrRNA2^13^. Disruption of the pseudoknot by mutation of either base (G3C and C40G) abolished the ability of the MVEV RNA to resist Xrn1 degradation^13^. Similarly, G3C mutant of ZIKV xrRNA1 also abolishes its Xrn1 resistance ability in our research (**Figure S9b**). SAXS data indicated that G3C mutant has larger *R_g_* and *D_max_* and Kratky plot with feature for partially folded molecules even in the presence of 5 mM Mg^2+^, as compared with ZIKV xrRNA1 (**Figure S10**, **Table S3**). These data suggest that G3C mutation could prevent PK1 formation and proper folding of xrRNA1 which results in disrupted ring structure. To study the significance of PK1 in ZIKV xrRNA1 mechanical anisotropy, we constructed both 5’_A36_-xrRNA1-G3C and 3’_A36_-xrRNA1-G3C which were used in nanopore sensing. For 5’_A36_-xrRNA1-G3C, the blockade current pattern was distinct from that of 5’_A36_-xrRNA1, the relative conductance was kept at a deep current state until RNA pass through the nanopore channel. As expected, the average duration time for 5’_A36_-xrRNA1-G3C (0.09 ± 0.04 s) decreased significantly with a reduction ratio of more than 1667 and 675 as compared with 5’_A36_-xrRNA1 (longer than 300 s) and 5’_A36_-xrRNA1-X (60.78 ± 5.32 s), respectively (**Figure 4a**). These data suggest that the PK1 pseudoknot contributes greatly to the 5’→3’ directional mechanical stability of xrRNA1. Nonetheless, both the blockade pattern and duration time of 3’_A36_-xrRNA1-G3C in the nanopore were similar to that of 3’_A36_-xrRNA1 (**Figure S11a**), therefore, PK1 pseudoknot interaction has minor effect on the 3’→5’ directional mechanical stability of xrRNA1, thus play an important role in the mechanical anisotropy of xrRNA1.

**Figure 4.**
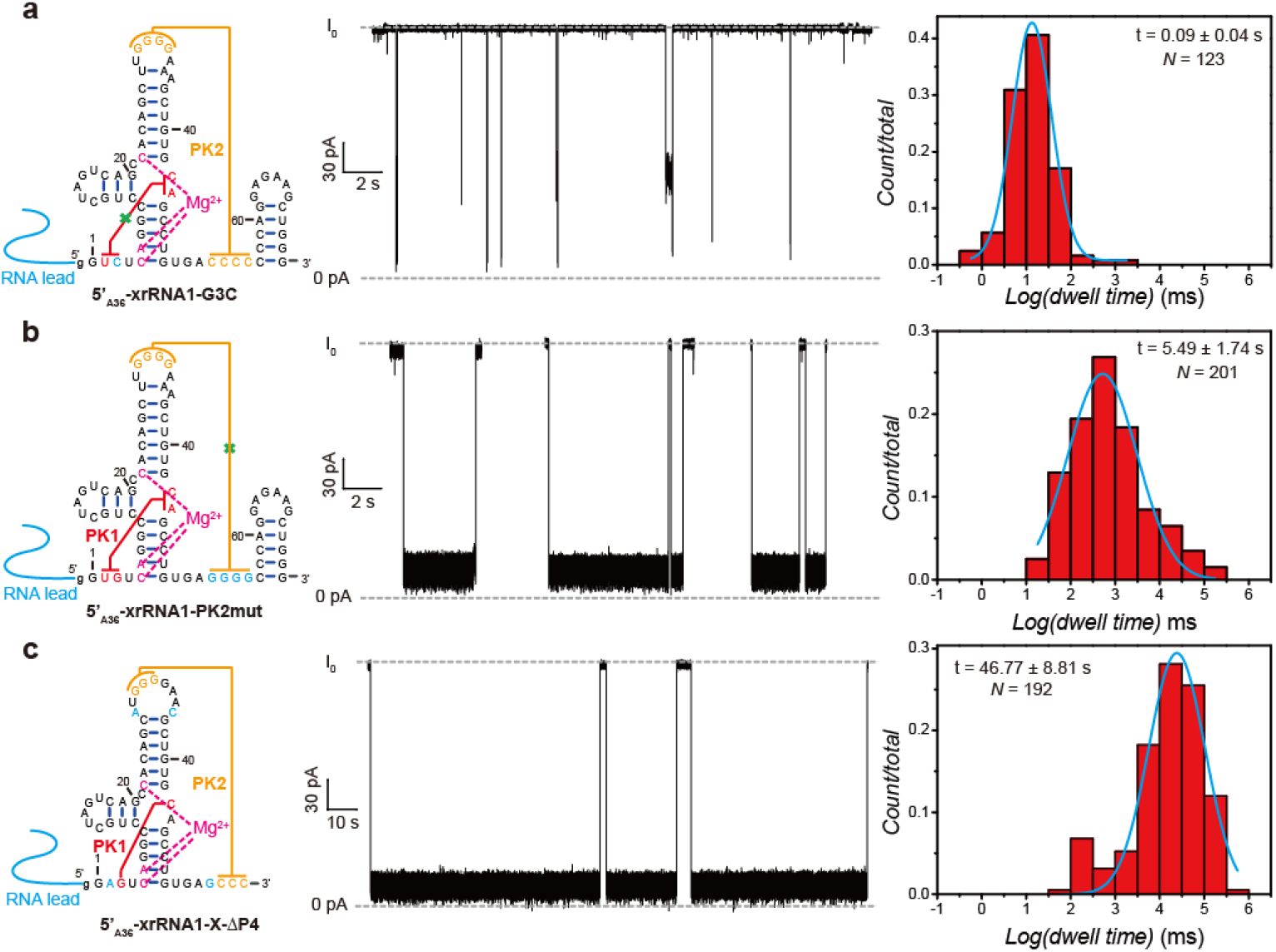
Contributions of key tertiary interactions to the directional mechanical stability of ZIKV xrRNA1. (**a-c**) Secondary structures (Left), representative current blockade traces (Middle) and the respective dwell time (Right) for the directional mechanical unfolding of 5’_A36_-xrRNA1-G3C (**a**), 5’_A36_-xrRNA1-PK2mut (**b**) and 5’_A36_-xrRNA1-X-ΔP4(**c**) in the α-HL nanopore.

The PK2 pseudoknot formed by base pairing between L3 and S4 is also crucial to the tertiary structure by latching the ring-like architecture, mutations that disrupt PK2 which 4Cs in S4 were replaced with 4Gs (xrRNA1-PK2mut) has been found to seriously weaken the resistance ability of ZIKV xrRNA1^14^. SAXS data indicate that xrRNA1-PK2mut has larger *R_g_* and *D_max_* as compared with ZIKV xrRNA1 and its Kratky plot has feature of partially folded molecule. The experimental scattering curve of xrRNA1-PK2mut fits poorly to ZIKV xrRNA1 crystal structure (**Figure S10a**). All these suggest the disruption of the ring structures. Similarly, 5’_A36_-xrRNA1-PK2mut and 3’_A36_-xrRNA1-PK2mut constructs were made to investigate the importance of the PK2 on the directional mechanical stability of ZIKV xrRNA1. While both constructs share similarities in blockade current signature as that of 5’_A36_-xrRNA1-X and 3’_A36_-xrRNA1-X, in which the current switches between two levels frequently, the elimination of the PK2 pseudoknot result in a shorter duration time (5.49 ± 1.74 s) for 5’_A36_-xrRNA1-PK2mut compared to 5’_A36_-xrRNA1 (>300 s) with reduction ratio of 53.6 (**Figure 4b**), but similar duration time (4.05 ± 0.54 s) for 3’_A36_-xrRNA1-PK2mut compared to 3’_A36_-xrRNA1 (2.06 ± 0.58 s). Hence, PK2 tertiary interaction also contributes significantly to the 5’→3’ but has marginally effects to the 3’→5’ directional mechanical stability of ZIKV xrRNA1, in accordance with our Xrn1 resistance assay that PK2-mut retains a weak resistance capability to Xrn1 while G3C totally can’t (**Figure S10e**).

To understand whether the coaxial stacking between the PK2 helix and P4 duplex contributes to the mechanical stability of ZIKV xrRNA1, we constructed P4 deletion mutant in the context of xrRNA1-X, which was referred as xrRNA1-X-ΔP4. The blockade current patterns for both 5’_A36_-xrRNA1-X-ΔP4 (**Figure 4c**) and 3’_A36_-xrRNA1-X-ΔP4 (**Figure S11c**) are similar to that of 5’_A36_-xrRNA1-X and 3’_A36_-xrRNA1-X, respectively. However, deletion of P4 slightly shortens the duration time of 5’_A36_-xrRNA1-X as compared with 5’_A36_-xrRNA1-X in the nanopore (46.77 ± 8.81 s vs 60.78 ±5.32 s) (**Figure 4c**). Thus, the coaxial stacking interaction between PK2 helix and P4 duplex contributes modestly to the 5’→3’ directional mechanical stability of xrRNA1. This result is consistent with our Xrn1 resistance assay that truncation of P4 marginally impairs ZIKV xrRNA1 ability to resist Xrn1 exoribonuclease, although the biological function of P4 is unknown (**Figure S10e**). SAXS analysis of xrRNA1-X-ΔP4 and xrRNA1-ΔP4 show that the experimental scattering profiles can nicely fit with the crystal structure of ZIKV xrRNA1 without P4, indicating that the truncation of P4 has little effect on the structural integrity of the ring-like core architecture of xrRNA1 (**Figure S10a**). The effect of truncation of P4 on the 3’ → 5’ directional mechanical stability was also probed. However, the duration time for 3’A36-xrRNA1-X-ΔP4 (4.2 ± 1.2 s) is only slightly longer than that of 3’_A36_-xrRNA1-X (2.0 ± 0.2 s) (**Table S4**).

### Mechanical anisotropy of ring-like RNA structures

Plotting the duration time of 5’ → 3’ and 3’ → 5’ directional translocation of ZIKV xrRNA1-X through the nanopore against Mg^2+^ concentrations clearly shows that the presence of high concentration of Mg^2+^ significantly promote the mechanical anisotropy in ZIKV xrRNA1 (**Figure 5a**), but the mechanical anisotropy is attenuated by disruption of the key tertiary interactions, even in the presence of Mg^2+^ (**Figure 5b**). Both Mg^2+^ and the tertiary interactions are important to facilitate xrRNA1 folding into its ring structure, suggesting that the mechanical anisotropy of ZIKV xrRNA1 correlates with the structural integrity of its ring-like topology structure.

**Figure 5.**
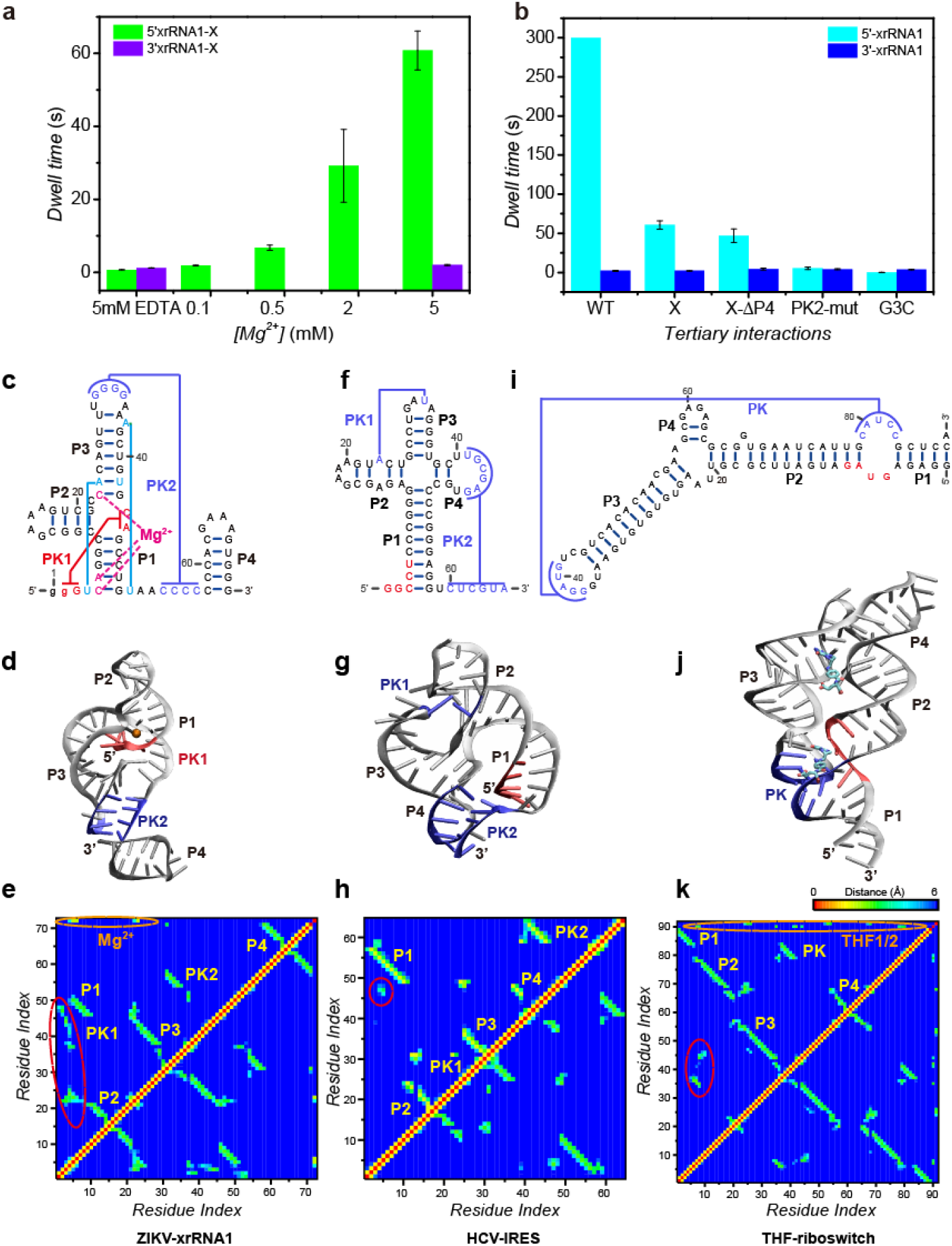
(**a-b**) Mechanical anisotropy of xrRNA1s is mainly dictated by its ring-like structure. (**a**) Plots of the dwell time for the unfolding and translocation of 5’_A36_-xrRNA1-X or 3’_A36_-xrRNA1-X in the α-HL nanopore against Mg^2+^ concentrations. (**b**) Plots of the dwell time for the unfolding and translocation of xrRNA1 mutants in the α-HL nanopore in the presence of 5 mM Mg^2+^. (**c-k**) Structural characteristics of the three representative ring-like RNAs. Secondary structures (the second row) and tertiary structures (the third row) of ZIKV-xrRNA1 (**c, f**), HCV-IRES (**d, g**) and THFriboswitch (**e, h**), respectively. The tightly bound magnesium ion or ligand are shown in sphere or stick representation, respectively. The pseudoknots enclosing the ring structures and the stretches passing through the rings are highlighted in blue, and red, respectively. (**i-k**) The residue-residue minimum distance map derived from crystal structure (upper left) and 100ns long MD simulation (lower right) for ZIKV-xrRNA1 (**i**), HCV-IRES (**j**) and THF-riboswitch (**k**), respectively. The tightly bound magnesium ion or ligand are also taken into consideration. Red circles represent the tertiary interaction of the 5’ end that passes through the ring, orange circles indicate the contacts that Mg^2+^ or THF ligand involved. Source data for panels **a** and **b** are provided as a Source Data File.

To further support the hypothesis and gain insights into the mechanisms underlying ZIKV xrRNA1 mechanical anisotropy, we used steered molecular dynamics simulation (SMD) to investigate the mechanical properties of the ZIKV xrRNA1 in addition to another two RNAs with ring-like structures, the core domain of the HCV IRES and the aptamer domain of THF riboswitch. HCV IRES is a highly structured RNA element within the 5’UTR of the HCV genome that mediates cap-independent translation initiation.^33^ As shown in **Figure 5f**, the core domain of the HCV IRES is organized as a four-way junction integrated with two pseudoknots. Unlike ZIKV xrRNA1, the 5’-terminal stretch (within P1) of the core domain of HCV IRES which threads through the ring almost has no tertiary contact with other parts of the RNA (**Figure 5d, e, g, h**). Tetrahydrofolate (THF) riboswitches are a group of RNA elements that bind tetrahydrofolate and are almost exclusively located in the 5’UTR of protein-coding genes involved in folate metabolism^34^. For THF riboswitch, the ring structure is enforced by a three-way junction (3WJ) motif between P2, P3 and P4, and a pseudoknot between L3 and J2/1^24^ (**Figure 5i**). The P1 stacks with the pseudoknot and lies outside of the ring, whereas a stretch between P1 and P2 (J1/2) passes through the ring instead. Two THF ligands which bind on 3-way junction and pseudoknot separately, could contribute to stabilization of the ring-like architecture^24^ (**Figure 5j, k**). In contrast to ZIKV xrRNA1, no cations are found to mediate the tertiary interactions in the crystal structures of two another ring structures. Additionally, the stretches passing through their ring structures are distal to the junction regions but close to the pseudoknots that enclose the ring, distinct from ZIKV-xrRNA1. Therefore, although these RNAs share with ring-like structural features, they still have considerable structural variations.

To model the unidirectional mechanical unfolding processes of the ring-like RNA structures within a nanocavity, a simple single-atom-layer nanopore with a diameter of ~13 Å which allows single-stranded RNA to thread through was employed in our simulations (**Figure 6a**). Such simple nanopore system avoids the putative complicated interactions between the pore and RNA, favorable to emphasize the intrinsic mechanical characteristics of RNA during translocation. We firstly studied the unfolding of ZIKV xrRNA1 induced by the 5’-end pulling at constant velocity of 0.1nm/ns (**Figure 6b, S12**). ZIKV xrRNA1 exhibits high mechanical stability against external force, and multiple force peaks are observed at the initial unfolding stage. The largest force peak height is up to 1600 pN, almost 3 times as high as that observed for the mechanical stable protein (i.e. ~550 pN for ubiquitin) using similar methods.^35^ The average resistant force over initial regime (from ZIKV xrRNA1’s threading into the nanopore to the position where highest force peak occurs) is larger than 800 pN. These initial unfolding events correspond to the rupture of PK1 as well as the breakage of magnesium coordination with the three phosphates of C5, A6 and C22. Similar unfolding behaviors can be observed in simulations at different pulling velocity of 1nm/ns and 0.01nm/ns (**Figure S12**). The extraordinary mechanical stability exhibiting in the initial regime upon 5’-end pulling could become a rate-limiting step for the whole translocation course of ZIKV xrRNA1, is therefore crucial to prevent ZIKV xrRNA1 from degradation by 5’-exoribonuclease (i.e. Xrn1). As compared to the mechanical stability upon 5’-end pulling, the mechanical stability to 3’-end pulling is much lower. The respective duplexes are sequentially unraveled from 3’-end in an order of P4-PK2-P1-P3, resulting in small force peaks which are no larger than 500 pN. Apparently, mechanical unfolding of ZIKV xrRNA1 in 3’ → 5’ direction is much easier than that from the opposite direction. For comparison, we also performed conventional two-end pulling simulations for ZIKV xrRNA1. As expected, unfolding of ZIKV xrRNA1 indeed starts from the weaker 3’-end, the unfolding pattern is similar to that observed in 3’-end translocation-coupled unfolding. Our SMD simulation results support the existence of significant mechanical anisotropy in ZIKV xrRNA1, consistent with the single-molecule nanopore experiments.

**Figure 6.**
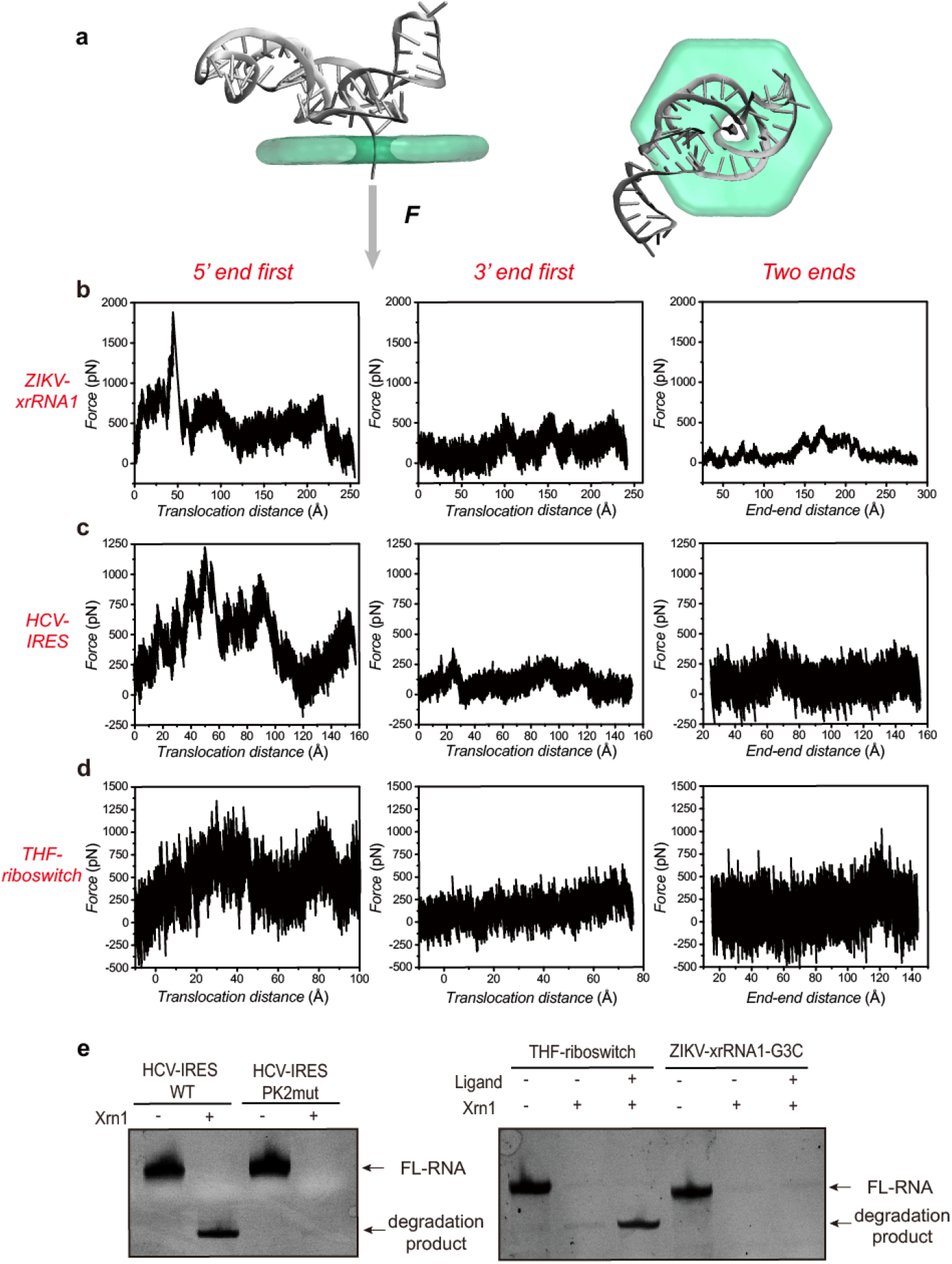
Mechanical unfolding of the three representative RNAs with ring-like topological structures using SMD simulations. (**a**) A schematic for the simulation system mimicking directional unfolding of the RNAs by nanopore. (**b-d**) Representative force profiles for mechanical unfolding of ZIKV-xrRNA1 (**b**), HCV-IRES (**c**) and THF-riboswitch (**d**) from different pulling directions (Left: 5’-end, Middle: 3’-end, Right: two-ends.) at a constant velocity of 0.1 nm/s. (**e**)Xrn1 resistance assay for RNAs. Source data for Panel **e** are provided as a Source Data File.

Significant mechanical anisotropy is also observed in the core domain of HCV IRES structure and THF-riboswitch in holo form (**Figure 6c, 6d and S13**). The high mechanical stability against 5’-end unfolding is further supported by the Xrn1-resistance assay (**Figure 6e**). For the HCV IRES, even no tertiary interactions exist between the 5’ end and other parts of the RNA, it still exhibits comparable mechanical resistance to 5’-end unfolding. As shown in **Figure 6e**, the disruption of the second pseudoknot in HCV IRES that enclose the ring totally abolish its capability to resist Xrn1 degradation. It’s likely the ring-like architecture plays a critical role in enhancing RNAs mechanical stability. In the case of THF riboswitch, its resistance to Xrn1 degradation disappears after removal of THF ligand (**Figure 6e**), which is crucial for THF riboswitch to maintain ring-like structure. Therefore, the mechanical anisotropy of THF riboswitch could be controlled by ligand binding.

Taken together, our SMD simulation results and biochemical assays on ZIKV xrRNA1, HCV IRES and the THF riboswitch support that the ring-like RNA structures could significantly enhance their mechanical anisotropy.

## DISCUSSION

In this study, we use the single-molecule nanopore sensing technique to investigate the directional mechanical stability of a viral RNA with ring-like structure, which the single-stranded leader sequences ensure unfolding-coupled translocation to be carried out in defined direction, therefore can better mimic the directionality of the cellular processes. We find that the mechanical stability of ZIKV xrRNA1 is dependent on the direction of the leader-guided translocation through the nanopore. In the presence of 5 mM Mg^2+^, the duration time of the translocation through the nanopore from the 5’ → 3’ direction is more than 150 times longer than that from the 3’ → 5’ direction, confirming extreme mechanical anisotropy in ZIKV xrRNA1. The extreme mechanical anisotropy is consistent with its functional activities in resisting degradation by cellular exonuclease Xrn1 to generate sfRNAs but readily unfolding as template for negative strand synthesis and genome replication induced by viral RdRP, therefore is of high biological significance. The nanopore sensing experiments show that the mechanical anisotropy of ZIKV xrRNA1 highly depends on Mg^2+^, which is a result of high dependence of 5’ → 3’ directional mechanical stability but no obvious dependence of 3’ → 5’ directional mechanical stability on Mg^2+^. The possessing of controllable mechanical stability and regulable mechanical anisotropy is important for biomolecules’ biomedical applications.

Mg^2+^ may facilitate xrRNA1 folding and structure and modulate the mechanical stability through both direct binding and acting as counterions to allow intramolecular tertiary interactions by reducing repulsive forces^36^. Crystal structure of ZIKV xrRNA1 has identified one Mg^2+^ ion binding site, mutations on the binding sites (C5G, A6U and C22G) results in loss of Xrn1 resistance ability, larger *R_g_* and *D_max_*, improper folding and increased flexibility even in the presence of 5 mM Mg^2+^ by SAXS (**Figure S14, Table S3**), supporting the importance of direct Mg^2+^ ion binding to xrRNA1 structure and folding. Mutational perturbation has also identified the importance of tertiary interactions such as PK1 and PK2 in xrRNA1 structure and mechanical stability, however, the formation and stabilization of such tertiary interactions must require Mg^2+^. In the absence of Mg^2+^, ZIKV xrRNA1 adopts unfolded structure and no such tertiary interactions are formed, supporting the critical roles of Mg^2+^ as counterions in shielding negative charges and reducing repulsive forces, therefore stabilizing such tertiary interactions. These results are also consistent with the DSC data which shows a more sensitive dependence of tertiary interactions on Mg^2+^ (**Figure S3**).

The ring-like architecture and the key tertiary interactions in xrRNAs are highly conserved across diverse mosquitoborne flaviviruses. The presence of duplicated xrRNA structures in the 3’ UTR is a common feature to most of the mosquito-borne flavivirus RNA genomes^37^. Recent research indicates that xrRNAs are also widespread in coding and noncoding subgenomic RNAs of two large families of plant-infecting RNA viruses^38^. The intricate mechanical anisotropy endowed by the ring-like architecture of xrRNAs may represent a more general strategy for RNA maturation and maintenance in many vectorial processes than previous known. Our nanopore sensing experiments also show that disruption of the key tertiary interactions in ZIKV xrRNA1 affects the 5’ → 3’ directional mechanical stability significantly, but has no obvious effects on the 3’ → 5’ directional mechanical stability, therefore, modulation of tertiary structure during infection such as mutations may mainly regulate sfRNA production but have minimal effect on flavivirus replication. A recent report showed that structural changes in the xrRNA structures of the dengue virus genome facilitates virus’s rapid adaptation to mosquito and human, moreover, adaptive mutations and deletions during host switch were mainly accumulated in regions that would effectively abrogate tertiary interactions^39^. It’s interesting to know how these mutations and deletions affect sfRNAs production in infected hosts, although having not been studied yet. sfRNAs production have been implicated to impact on flavivirus replication, cytopathicity and pathogenicity^10, 11, 40, 41^. We speculate that sfRNAs production modulated by the tertiary structural changes upon host switch is related to the viruses’ ability to adapt to different hosts.

Understanding the mechanisms that determine RNA mechanical anisotropy could be of great importance in development of RNA-based mechanically anisotropic biomaterials. Due to its diverse structural and functional attributes, RNA has recently attracted widespread attention as a unique biomaterial with distinct biophysical properties for designing sophisticated architectures in the nanometer scale.^42^ For example, the φ29 pRNA three-way-junction (3WJ) has been used as building block to construct RNA triangles, squares, pentagons, and hexagons, as a platform for building a variety of multifunctional nanoparticles as potential therapeutic agents^42, 43, 44^. Mechanical anisotropy is an essential property for many soft biological tissues. The microstructures of many such tissues, such as the fibers of the human Achilles tendon, often run parallel to each other in a specific direction, leading to anisotropy in the mechanical properties of the tissues, therefore have been identified as typical anisotropic materials^45^. However, it remains a big challenge to build biomimetic materials with mechanical anisotropy based on macromolecules. Discovering molecular building blocks with anisotropic mechanical properties and revealing the structural mechanisms of mechanical anisotropy is the key step to overcome this challenge. Our work shows that the mechanical anisotropy of a flaviviral RNA is largely defined by the structural integrity of its ring-like topological structure, which is regulated by Mg^2+^ and the key tertiary interactions that act cooperatively to facilitate RNA folding. Mechanical anisotropy is also observed in another two RNAs with ring-like structures by SMD, suggesting the generality of the relationship. Interestingly, the mechanical anisotropy of ZIKV xrRNA1 and the aptamer of THF riboswitch can be reversibly regulated by Mg^2+^ and ligand binding. Most of the mechanically anisotropic biomaterials cannot respond to external stimuli^46^. Therefore, both ZIKV xrRNA1 and the aptamer of THF riboswitch could be ideal building blocks for environmental responsive anisotropic artificial tissues. It’s expected that more ring-like RNA structures with mechanical anisotropy could be identified and designed for development of RNA-based mechanically anisotropic biomaterials.

## METHODS

### RNA sample preparation

The wild type ZIKV xrRNA1 and its mutant constructs were generated as follows. Plasmids coding an upstream T7 promoter, the ZIKV xrRNA1 sequence with its upstream or downstream polyA sequences, were gene synthesized and sequenced by Wuxi Qinglan Biotechnology Inc, Wuxi, China. All the other mutants were generated using the Transgen’s Fast Mutagenesis System. The double-stranded DNA fragment templates for in vitro RNA production were generated by PCR using an upstream forward primer targeted the plasmids and a downstream reverse primer specific to respective cDNAs. The RNAs were transcribed in vitro using T7 RNA polymerase and purified by preparative, non-denaturing polyacrylamide gel electrophoresis, the target RNA bands were cut and passively eluted from gel slices into buffer containing 0.3 M NaOAc, 1 mM EDTA, pH 5.2 overnight at 4 ℃. The RNAs were further passed through the size exclusion chromatography column to final buffer condition for nanopore sensing, DSC, SAXS and other biochemical experiments. The sequences for all the constructs used in this study are listed in **Table S5**.

### Differential scanning calorimetry

All DSC measurements were performed using a VP DSC Micro-calorimeter from the Malvern MicroCal (Northampton, MA). The DSC consists of a matched pair of 0.511 ml sample and reference cells. In a series of DSC scans, both cells were first loaded with the buffer solution, equilibrated at 20 °C for 15 min, and scanned from 20 to 100°C at a scan rate of 100 °C /h. The buffer versus buffer scan was repeated five times to obtain baseline, then the sample cell was emptied, rinsed, and loaded with the RNAs solution prior to the 15 min equilibration period. RNA samples were kept at a concentration of 30 μM and in buffers containing 10 mM sodium phosphate, 150 mM KCl, pH 7.5 and various concentrations of Mg^2+^ and care was taken to minimize the presence of air bubbles in loading of the sample cell. The experimental data is processed by subtracting the baseline firstly, then normalizing the concentration and using a non-two-state model to deconvolute the curve by using MicroCal software, finally the melting temperature *Tm* which relates to stability was obtained directly^47, 48^.

### Small angle X-ray scattering

All the parameters for data collection and software employed for data analysis are summarize in **Table S6**. SAXS measurements were carried out at room temperature at the beamline 12 ID-B of the Advanced Photon Source, Argonne National Laboratory or the beamline BL19U2 of the National Center for Protein Science Shanghai (NCPSS) and Shanghai Synchrotron Radiation Facility (SSRF). The scattered X-ray photons were recorded with a PILATUS 2M detector (Dectris) at 12 ID-B and a PILATUS 100k detector (Dectris) at BL19U2. The setups were adjusted to achieve scattering *q* values of 0.005 <*q*< 0.89 Å^−1^ (12ID-B) or 0.009<*q*< 0.415Å^−1^ (BL19U2), where *q* = (4π/λ)sinθ, and 2θ is the scattering angle. Thirty two-dimensional images were recorded for each buffer or sample solution using a flow cell, with the exposure time of 0.5-2 seconds to minimize radiation damage and obtain good signal-to-noise ratio. No radiation damage was observed as confirmed by the absence of systematic signal changes in sequentially collected X-ray scattering images. The 2D images were reduced to one-dimensional scattering profiles using Matlab (12ID-B) or BioXTAS Raw (BL19U2). Scattering profiles of the RNAs were calculated by subtracting the background buffer contribution from the sample-buffer profile using the program PRIMUS^49^ following standard procedures. Concentration series measurements (4- and 2-fold dilution and stock solution) for the same sample were carried out to remove the scattering contribution due to inter-particle interactions and to extrapolate the data to infinite dilution. The forward scattering intensity *I*(0) and the radius of gyration (*R*_g_) were calculated from the data of infinite dilution at low *q* values in the range of *qR*_g_< 1.3, using the Guinier approximation: ln*I*(*q*)≈ln(*I*(0))-*R*_*g*_^2^*q*^2^/3. These parameters were also estimated from the scattering profile with a broader *q* range of 0.006-0.30 Å^−1^ using the indirect Fourier transform method implemented in the program GNOM^50^, along with the pair distance distribution function (PDDF), *p*(*r*), and the maximum dimension of the protein, *D*_max_. The parameter *D*_max_ (the upper end of distance *r*), was chosen so that the resulting PDDF has a short, near zero-value tail to avoid underestimation of the molecular dimension and consequent distortion in low resolution structural reconstruction. The Volume-of-correlation (*V*_c_) were calculated using the program Scatter, and the molecular weights of solutes were calculated on a relative scale using the *R*_g_/*V*_c_ power law developed by Rambo *et al*^51^, independently of RNA concentration and with minimal user bias. The theoretical scattering intensity of the atomic structure model was calculated and fitted to the experimental scattering intensity using CRYSOL^52^.

### Xrn1 resistance assay

The Xrn1 resistance experiments were conducted by following the standard protocol developed in previous studies^53^. Recombinant Xrn1 from *Kluyveromyces lactis* and RppH from *Bdellovibrio bacteriovorus* were expressed in *E.coli* and purified by Ni^2+^-NTA affinity and size-exclusion chromatography. In a 10ul reaction, ~600 ng purified RNA was treated with 0.5ul of >3 U/μl RppH and then split between two tubes. 0.5ul of >3 U/μlXrn1 was added to one half of the reaction while the other served as a (−) Xrn1 control, followed by incubation at 37℃ for 30min. Reactions were then quenched by the addition of an equal volume of a TBE-Urea loading dye containing 30 mM EDTA, 8 M Urea and 0.1% (wt/vol) xylene cyanol and bromophenol blue. Then the RNA products were analyzed on 8% TBE-Urea denaturing gel and visualized by staining with Gel safe.

### Single-molecule nanopore sensing and data analysis

Single-channel current recordings were performed with an individual α-hemolysin nanopore inserted into a vertical lipid bilayer. The vertical chamber setup was assembled by two compartments, a cuvette with a 150 μm aperture drilled on the side (*cis*) and a bilayer chamber (*trans*) (Warner Instruments, Hamden, CT, USA). After pre-painting both sides of the cuvette aperture with 0.5 mg/mL DPhPC/hexane, both chambers were filled with 1 mL of test buffer (20 mM Tris, pH 7.5, 1M KCl (*cis*)/3M KCl (*trans*) with different Mg^2+^ concentrations). RNAs were added to the cis solution with a final concentration of 50 nM. The lipid bilayer was created by applying 30 mg/mL DPhPC/decane to the pretreated aperture. *In vitro* assembled WT-α-HL nanopore protein was added in the *cis* compartment. After a single protein nanopore was formed in the lipid bilayer, positive potentials of 100 mV were applied across the lipid bilayer with Ag/AgCl electrodes. The cis compartment was defined as the virtual ground. The electrical current was recorded with a patch-clamp amplifier (HEKA EPC10; HEKA Elektronik, Lambrecht/Pfalz, Germany). Recordings were collected using a 3 kHz low-pass Bessel filter at sampling frequency of 20 kHz with a computer equipped with a LIH 1600 A/D converter (HEKA Elektronik). All measurements were carried out at room temperature. Data analysis was performed using MATLAB (R2011b, MathWorks) software and OriginLab 8.0 (OriginLab Corp., Northampton, MA, USA). The current blockades are described as *I*/*I*_0_, where *I*_0_ is the ionic current of open nanopore and *I* is the blockage current produced by the analyte. Events with current blockades larger than 70% and dwell time longer than 0.15 ms were analyzed as RNA translocations. The mean dwell time for current spikes was obtained from the dwell time histograms. All the data are presented as mean ± standard error of the mean of three independent experiments.

### Molecular dynamics simulations

In the spirit of nanopore sensing experiments, we perform SMD simulations in a artificial nanopore system, which was modeled by one graphene layer with a 13 Å diameter pore in its center and is comparable to the pore size of α-hemolysin^54^. Three representative RNAs with ring-like architecture including flaviviral Xrn1-resistant RNA (ZIKV xrRNA1, PDB code: 5TPY), the core domain of Hepatitis C virus internal ribosome entry site (IRES, PDB code: 3T4B), and the aptamer domain of THF-riboswitch in holo form (PDB code: 4LVV), were selected to study their mechanical properties^23, 24, 55^. Those RNAs show diverse topologies and biological functions. For core structure of HCV-IRES, the tetraloop receptor which acts as crystallization module was removed in our studies, to better characterize the mechanical properties of the ring-like structure. These RNAs were subjected to equilibrium simulations in an isolated system, following the similar protocol in the previous study^56^. After 100ns equilibrium simulations without any restraint, the equilibriumed RNA configuration along with the water in first hydration shell (the cutoff distance is 3.5 Å), was extracted to be used as a starting structure for subsequent simulations. Then the RNA structure was translated and rotated so that the pulling terminus (5’ or 3’ terminus) was above the pore. The merged system was further equilibriumed within NPT ensemble for 10 ns with positional restraint on the pulling terminus, allowing RNA to further relaxation and free rotation around the anchor. Then the external force was loaded on the chosen terminus of RNA using Steered Molecular Dynamics (SMD) simulation, to make the RNA thread through the nanopore. The position restraints were imposed on the nanopore, to mimic immobilization of the pore. For comparison, we also carried out conventional two-end pulling simulations for each RNA.

All simulations were conducted using GROMACS 2016 or 2018 packages^57, 58^.The parameters for the RNA and monovalent ions were taken from the AMBER14 Force Field and used in concert with the TIP3P explicit water model^59, 60, 61, 62^. The Mg^2+^ ion model parameterized by Villa et al was used in simulations^63^.The parameters for THF were taken from General Amber Force Field (gaff)^64^ combined with Restraint Electrostatic Potential Charge (RESP), both of which are compatible to AMBER Force Field. The parameters for nanopore were adapted from the aromatic carbon atom in AMBER Force Field (e.g CA), and the atomic charge is set to zero as previous computation works^35, 65^. To avoid the putative sticking of aromatic ring (i.e. RNA base and THF) on nanopore surface, the interaction strength (∊, the depth of Lennard-Jones potential) between nanopore and RNA/THF were reduced to one tenth of the original. Monovalent ions (K^+^ and Cl^−^) were added to neutralize the system, yielding an ionic concentration of 150 mM. Periodic boundary conditions were applied to all the systems in three directions. van der Waals and short-range electrostatic interactions were calculated at a cut-off distance of 10 Å, while long range electrostatic interactions were computed by the particle mesh Ewald (PME) method^66^. The temperature (*T* = 310 K) and pressure (*P* = 1 atm) were maintained using a stochastic velocity rescaling thermostat and Parrinello-Rahman barostat, respectively. All SMD simulations were run in NVT ensemble at a temperature of 310 K. Partial structural visualization and analysis were performed using VMD1.93^67^.

## ACKNOWLEDGMENT

We thank Dr. Xiaobing Zuo at the beamline 12-ID-B, Advanced Photon Source, Argonne National Laboratory and the staffs of beamline BL19U2 at Shanghai Synchrotron Radiation Facility, Shanghai, China for assistance during SAXS data collection.

