## Supplementary material for "Molecular mechanisms underlying the extreme mechanical anisotropy of the flaviviral exoribonuclease-resistant RNAs (xrRNAs)": SI-xrRNAMechanicalStability-BioRxiv.pdf

### EXTENDED SECTIONS

This section is intended to provide an elaborate discussion of certain results of the main text that had to keep brief because of length restrictions.

**$Mg^{2+}$-dependent structural transition and thermal stability of ZIKV xrRNA1.** The formation of stable and compact RNA structure requires the neutralization of polyanionic phosphate backbone by counterions[1]. An important question is the extent to which tertiary interactions stabilize the ZIKV xrRNA1 ring structure in the presence of different types of counterions, such as $K^+$, $Na^+$, $Mg^{2+}$, etc. $Mg^{2+}$ ions are known to be important for the structure and stability of RNA molecules[2]. As shown in **Figure 1a-b**, a $Mg^{2+}$ ion is found to coordinate with the phosphates of C5, A6 and C22 in the crystal structure of ZIKV xrRNA1, implying that $Mg^{2+}$ ions may play an important role in the structure and folding of xrRNA1.

To understand how $Mg^{2+}$ affect the overall structure and folding of ZIKV xrRNA1, small angle x-ray scattering (SAXS) experiments were carried out to follow the $Mg^{2+}$-induced structural transition. The representative scattering profiles, with scattering intensity $I(q)$ plotted against momentum transfer $q$, the Guinier fitting of the scattering profiles, along with pair distance distribution function PDDFs transformed from scattering profiles for ZIKV xrRNA1 in 20 mM Tris-HCl supplemented with different concentration of $Mg^{2+}$, are shown in Figure S2a-c. The overall structural parameters, including $R_g$ calculated from Guinier slops, $R_g$ and $D_{max}$ from the PDDF functions, as well as molecular weights derived from volume-of-correlation ($V_c$),[3] are summarized in **Table S1**. Plotting the radius of gyration ($R_g$) values calculated from the PDDF functions against $Mg^{2+}$ concentrations shows a cooperative change in $R_g$ value, suggesting that ZIKV xrRNA1 undergoes a $Mg^{2+}$-induced two-state transition between the unfolded and folded states, and the midpoint of the transition is about 0.3 mM $Mg^{2+}$ (**Figure S2e**). The fraction of unfolded RNA $\Phi_U$ was calculated at each $Mg^{2+}$ concentration from $R^2_g = \Phi_U R^2_{g,U} + (1-\Phi_U)R^2_{g,F}$, with $R_{g,U}$ and $R_{g,F}$ being the radii of the gyration for the unfolded and folded RNA in 20 mM Tris-HCl and 5 mM $Mg^{2+}$, respectively (**Figure S2f**). The transition from an open, unfolded conformation to a globular, folded form is also apparent from the changes of Kratky plots of xrRNA1 in 0 to 10 mM $Mg^{2+}$ (**Figure S2d**). In 20 mM Tris-HCl and the absence of $Mg^{2+}$, ZIKV xrRNA1 may form secondary structures such as duplexes but no discernible high-order structure, therefore ZIKV xrRNA1 is in an open, unfolded state characterized with an average $R_g$ of ~30Å. Upon increasing the $Mg^{2+}$ concentration, ZIKV xrRNA1 becomes more compact, and the $R_g$ shrink to ~23 Å at $Mg^{2+}$ concentrations above ~1mM and doesn't change significantly between 2 to 10 mM $Mg^{2+}$, therefore ZIKV xrRNA1 mainly adopts a folded conformation with the tertiary structures formed, which is further supported by the fact that the experimental scattering profile at 5 mM $Mg^{2+}$ fits nicely to the theoretical scattering curve calculated from the crystal structure.[4] The effect of monovalent cation, such as $K^+$ on the overall structure of ZIKV xrRNA1 is also probed. In comparison with that in 20 mM Tris-HCl and no $Mg^{2+}$, the average $R_g$ value of xrRNA1 at 20 mM Tris-HCl, 100 mM $K^+$ deceases to ~24.45Å, but doesn't change significantly upon even higher concentration of $K^+$ such as at 20 mM Tris-HCl, 1000 mM $K^+$ ($R_g$=23.47Å) (**Table S1**). These results suggest that $Mg^{2+}$ is generally a more potent stabilizer of the tertiary interactions than $K^+$ ions and essential to the overall structure of ZIKV xrRNA1.

The effects of $Mg^{2+}$ on the overall thermal stability of ZIKV xrRNA1 was analyzed by differential scanning calorimetry (DSC) which can provide RNA stability in terms of melting temperature.[5] The DSC thermograms of ZIKV xrRNA1 are compared over a range of $Mg^{2+}$ concentrations (**Figure S3a**). **Figure S3b** shows a typical DSC thermogram for ZIKV xrRNA1 in 0 mM $Mg^{2+}$, which can be further fitted into two distinct transition peaks with $T_m$ values of 63.3, 75.3 °C, respectively. The low- and high- temperature transitions can be attributed to unfolding of tertiary and secondary structures, respectively.[6] Both the low- and high- temperature transitions shift to high temperature as $Mg^{2+}$ concentration increases, but the low- is more sensitive than the high- temperature transition to changes in $Mg^{2+}$ concentrations, resulting in merging of these two transitions into one peak above 5 mM $Mg^{2+}$ (**Figure S3c**) which xrRNA1 is thought to unfold in a cooperative manner. These results suggest an essential role of $Mg^{2+}$ in stabilizing the tertiary interactions rather than the secondary structure in ZIKV xrRNA1.

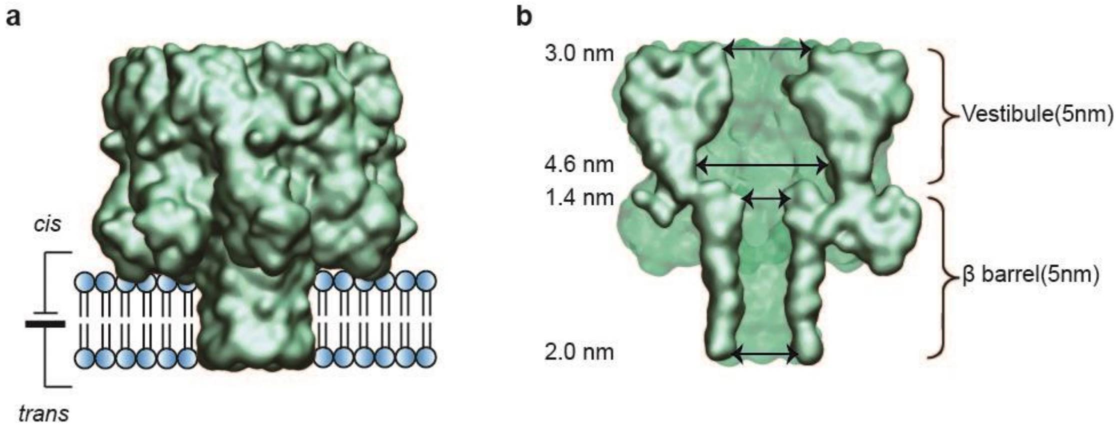

**Figure S1**. Dimensions of the α-hemolysin nanopore. (**a**) Space filling model showing the α-HL nanopore inserted in the phospholipid bilayer. (**b**) Cross section view and dimensions of the α-HL nanopore.

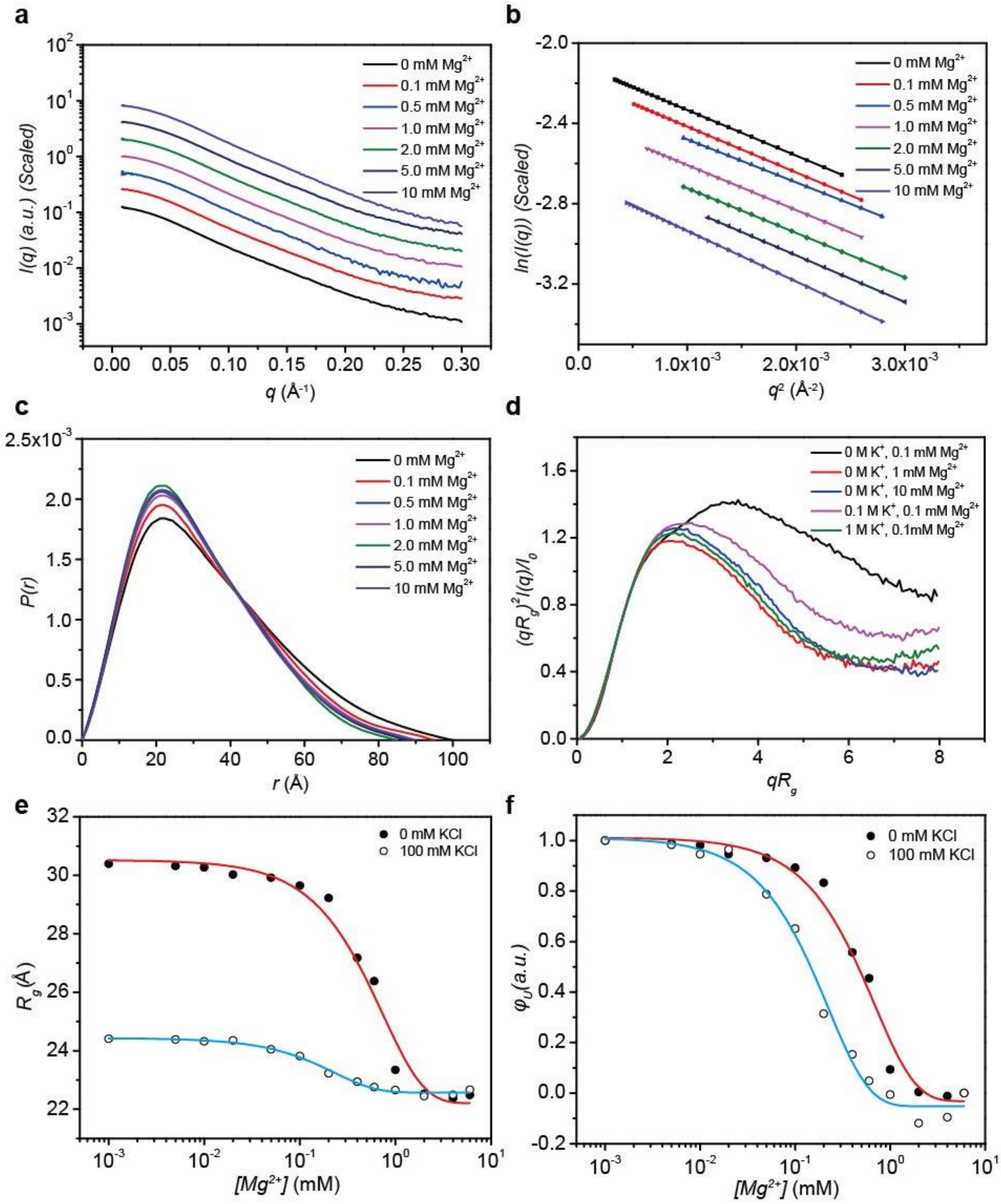

**Figure S2**. $Mg^{2+}$-dependent structural transition of ZIKV xrRNA1. (**a-d**) Experimental scattering profiles (**a**), guinier fitting (**b**), PDDFs and dimensionless Kratky plots (**d**) for ZIKV-xrRNA1 in 20 mM Tris-HCl, pH 7.50 supplemented with different $Mg^{2+}$ concentrations. (e) Plots of $R_g$ as a function of [$Mg^{2+}$] in buffers of 20 mM Tris-HCl (black solid circle), pH 7.50, or 20 mM Tris-HCl, 100 mM $K^+$, pH 7.50 (cyan open circle), where $R_g$ was obtained from PDDF functions of SAXS data. (**f**) Fraction of unfolded RNA $\Phi_U$ versus [$Mg^{2+}$]. Symbols are as for (**a**).

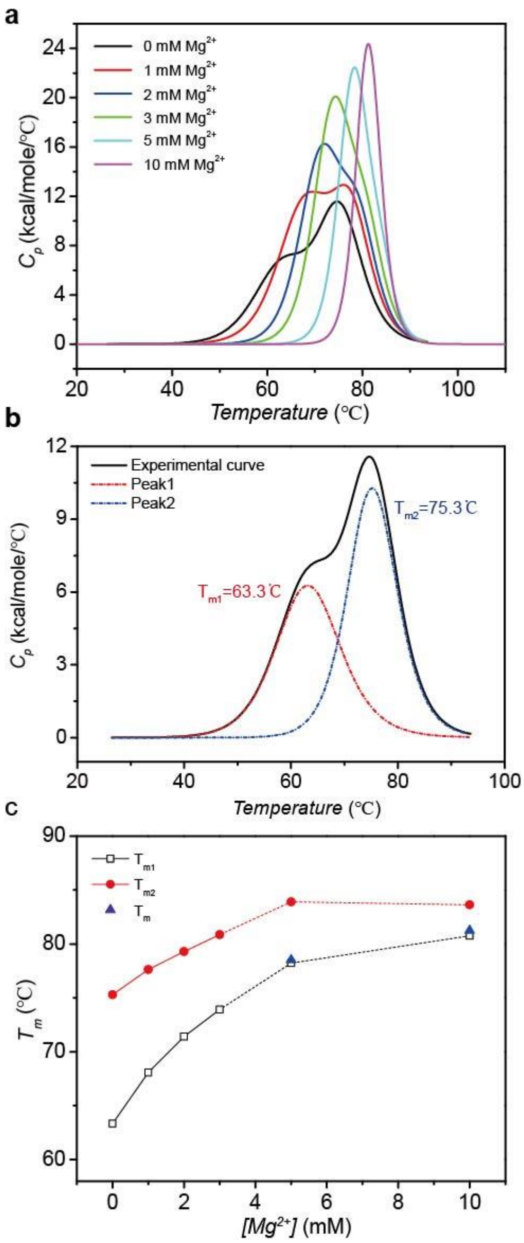

**Figure S3**. DSC analysis of the thermal stability of ZIKV-xrRNA1. (**a**) Thermal unfolding of ZIKV-xrRNA1 samples at a concentration of 30 μM and in buffers containing 10 mM sodium phosphate, 150 mM KCl, pH 7.5 and various concentrations of $Mg^{2+}$ recorded by DSC. (b) A typical DSC thermogram for ZIKV-xrRNA1 in 0 mM $Mg^{2+}$. DSC thermograms at low $Mg^{2+}$ concentrations (< 5mM) can be fitted into two distinct transition peaks with two $T_m$ values $T_{m1}$ and $T_{m2}$, corresponding to the unfolding of tertiary and secondary structure respectively, whereas with the increasing of $Mg^{2+}$ concentrations, DSC thermograms for xrRNA1 can only be fitted into one peak with one transition temperature $T_m$, which xrRNA1 is considered to unfold cooperatively. (**c**) $T_{m1}$, $T_{m2}$ and $T_m$ values extracted from DSC were plotted against concentration of $Mg^{2+}$.

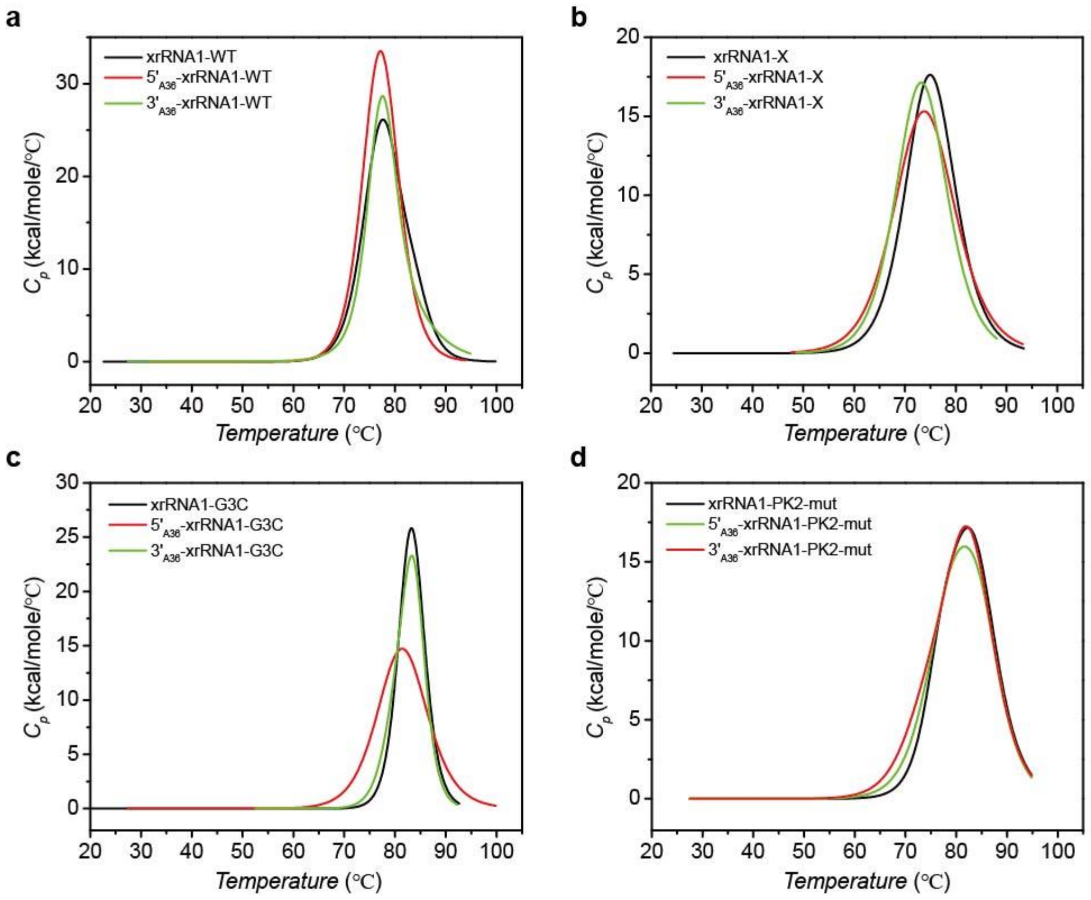

**Figure S4**. The effects of the 5' or 3' poly(rA) leader sequences on the thermal stability of ZIKV xrRNA1 and its mutants. (**a-d**) The DSC thermograms for ZIKV xrRNA1 (**a**), xrRNA1-X (**b**), xrRNA1-G3C (**c**) and xrRNA1-PK2mut (**d**) and their related constructs with 5' or 3' leader sequences show that the 5' or 3' leader sequences have minor effects on thermal stability of the corresponding RNAs. All the experiments were conducted in buffers containing 5 mM $Mg^{2+}$.

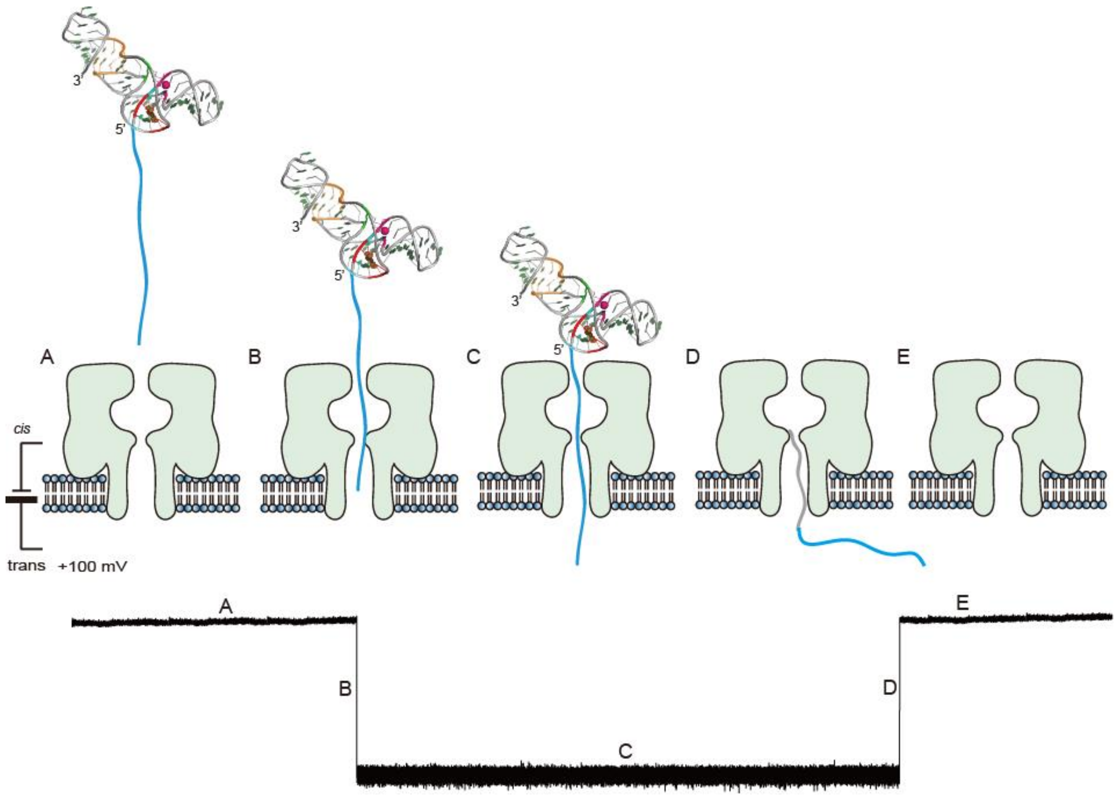

**Figure S5**. Schematic representation of xrRNA1 translocation and unfolding in the α-HL nanopore. A. open pore; B. leader sequence enters into the pore; C and D. RNA unfolds and then release; E. open pore. Due to the complexity of RNA structure and limitation of nanopore detection, it's difficult to have a thorough molecular interpretation of the blockade current traces but the duration time that RNA is trapped in the pore provides a measure of the RNAs' mechanical stability in a defined direction.

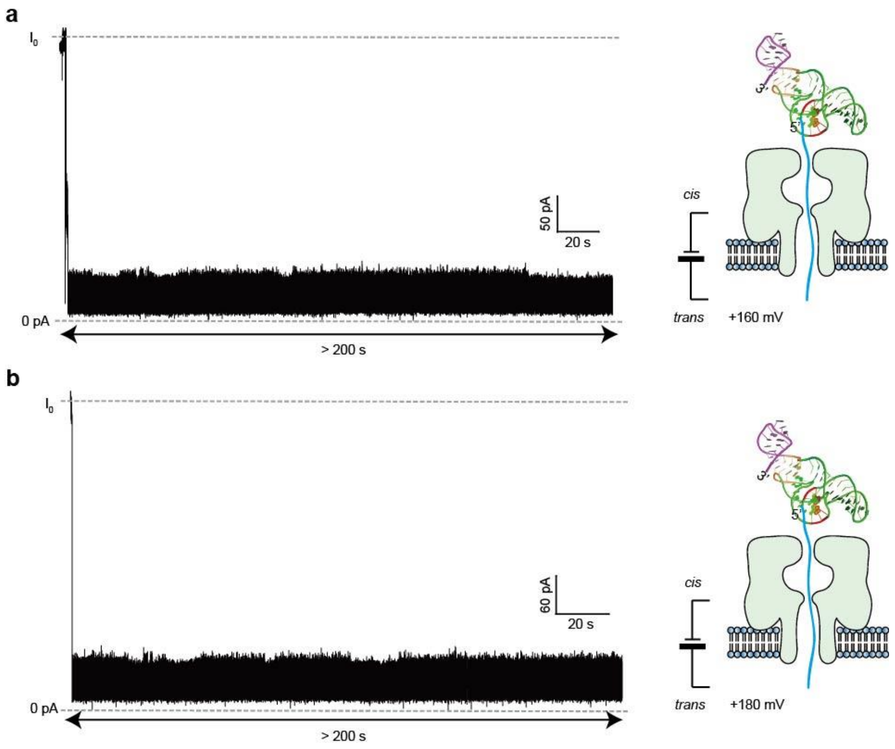

**Figure S6**. Typical current blockade traces for 5'$_{A36}$-xrRNA1 in 5 mM $Mg^{2+}$ in the α-HL nanopore under 160 mV (**a**) and 180 mV (**b**) transmembrane voltages.

a

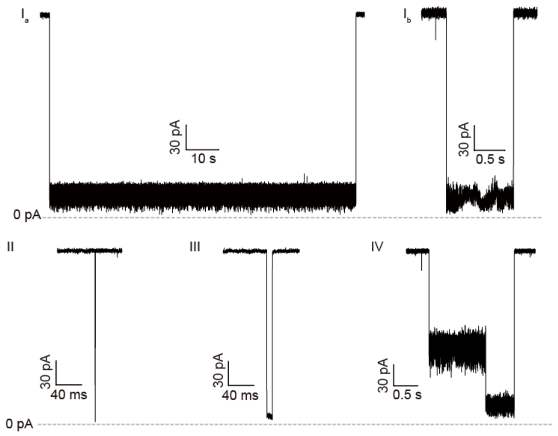

b

| Type | $3'_{A36}$-xrRNA1 | $5'_{A36}$-xrRNA1-X | $3'_{A36}$-xrRNA1-X | $5'_{A36}$-xrRNA1-G3C | $3'_{A36}$-xrRNA1-G3C | $5'_{A36}$-xrRNA1-PK2mut | $3'_{A36}$-xrRNA1-PK2mut | $5'_{A36}$-xrRNA1-X-ΔP4 | $3'_{A36}$-xrRNA1-X-ΔP4 |
|---|---|---|---|---|---|---|---|---|---|
| I | 85% | 70.91% | 79% | 1% | 89% | 75% | 84% | 71% | 72% |
| II | 5% | 13.64% | 5% | 10% | 2% | 4% | 6% | 11% | 5% |
| III | 4% | 11.82% | 11% | 51% | 3% | 17% | 9% | 18% | 13% |
| IV | 6% | 3.64% | 5% | 38% | 6% | 4% | 1% | 0 | 10% |

c

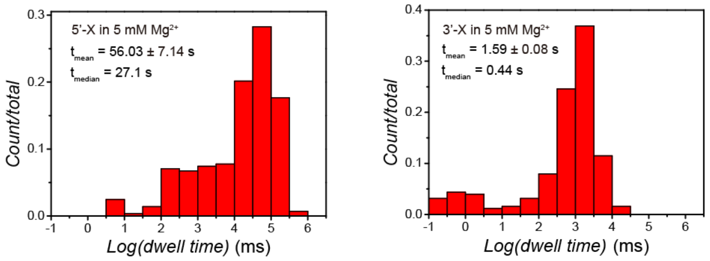

| mean dwell time | 5'X | 3'X | median dwell time | 5'X | 3'X |
|---|---|---|---|---|---|
| type 1 | 60.78 ± 5.32 s | 2.00 ± 0.15 s | type 1 | 35.81 s | 1.40 s |
| all signal | 56.03 ± 7.14 s | 1.59 ± 0.08 s | all signal | 27.10 s | 0.44 s |

**Figure S7**. Nanopore signatures for different RNA constructs in this study. (**a**) Typical current blockade signatures generated by $5'_{A36}$-xrRNA1-X and $3'_{A36}$-xrRNA1-X in the presence of 5 mM $Mg^{2+}$. (**b**) Populations of different current blockade types generated by xrRNA1 and its mutants. (**c**) Dwell time distribution for $5'_{A36}$-xrRNA1-X and $3'_{A36}$-xrRNA1-X and the statistical analysis was based on all the signal collected not the type1 only as in **Figure 3c**. The mean duration time for $5'_{A36}$-xrRNA1-X is ~ 35 folds longer than that for $3'_{A36}$-xrRNA1-X. Type1 represents the most typical events and is attributed to 3'→5' directional translocation guided by the leader sequence. Type 2 is characterized with mean duration time less than 1 ms, which could be the translocation signals of single-stranded RNAs due to the degradation or unfolding of xrRNA1. Type 3 typically show simple deep blockade current level ($\%I/I_0$ = 10%) and short duration time (~0.16s), which should be the signals for partially folded xrRNA1. Unlike the other three types, type 4 has two distinct sub-states. The ionic current initially drops to an intermediate current level of $\%I/I_0$ = 40%, indicating that not the narrow constriction zone of the β-barrel but the cis vestibule of the pore is occupied at this stage, then is dominated by the deep blockade current level of $\%I/I_0$ = 10% for long before re-establishing the open-pore current. Type 2-4 do not feature the directional unfolding and translocation guided by the leader sequence.

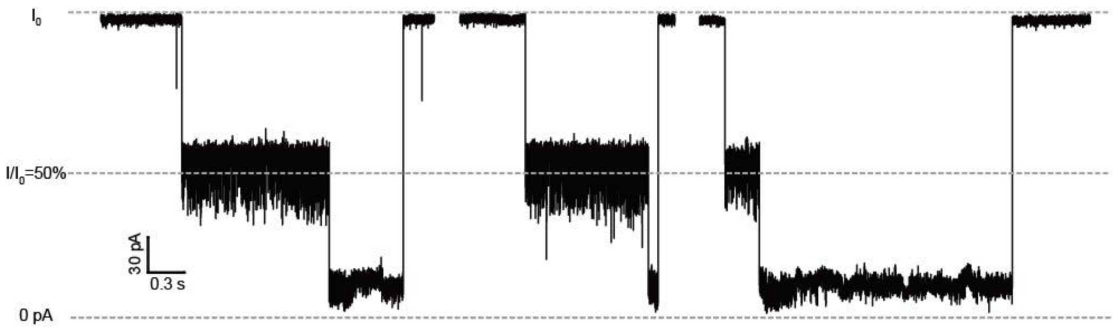

**Figure S8**. Nanopore current blockade signatures generated by ZIKV xrRNA1 without a leader sequence in buffer containing 5 mM $Mg^{2+}$. xrRNA1 generated a two-level blockade current pattern that the current firstly reduced to 50% of $I_0$, then reduced to a deep state with $I/I_0$= 10%. Previous study showed that DNA G-quadruplex is able to enter into the cis vestibule of the nanopore and cause blockade current level of $I/I_0$= 50%[7]. Given the structure of ZIKV xrRNA1 and the pattern of blockade current generated, it's likely that the L3-S4 pseudoknot (PK2) is conformationally dynamic or transiently unformed, the P4 helix at the 3' end of ZIKV xrRNA1 can be trapped in the vestibule of the nanopore first, then guide the unfolding and translocation of the remaining part of ZIKV xrRNA1 through the nanopore. Evidence for the dynamics of PK2 has been observed in the crystal structure of MVEV xrRNA2[8], which the PK2 is absent but highly homologous to ZIKV xrRNA1.

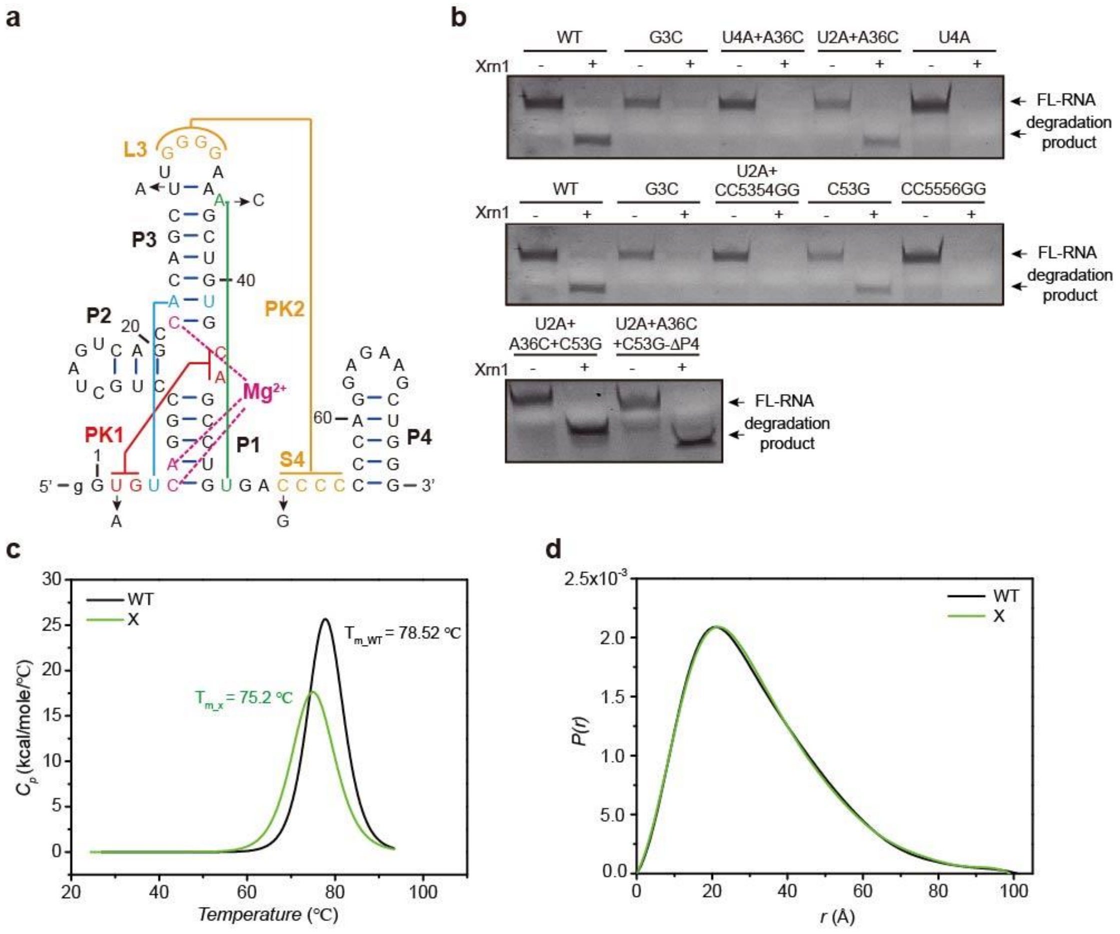

**Figure S9**. Screening a quadruple mutant xrRNA1-X to resemble xrRNA1. (**a**) Secondary structure of ZIKV xrRNA1, and the mutation site for a quadruple mutant xrRNA1-X are indicated with arrows. (**b**) Xrn1 resistance assay for ZIKV xrRNA1 and its various mutants. All the mutants were co-transcribed with a 36-nt poly(rA) leader sequence in the 5'end. (**c**) DSC thermograms for ZIKV xrRNA1 and xrRNA1-X in 5 mM $Mg^{2+}$. (**d**) The PDDF of xrRNA1-X can be nicely superimposed on that of ZIKV xrRNA1 in 5 mM $Mg^{2+}$. xrRNA1-X shows similar structure and thermal stability as ZIKV xrRNA1. Source data for panel **b** are provided as a Source Data File.

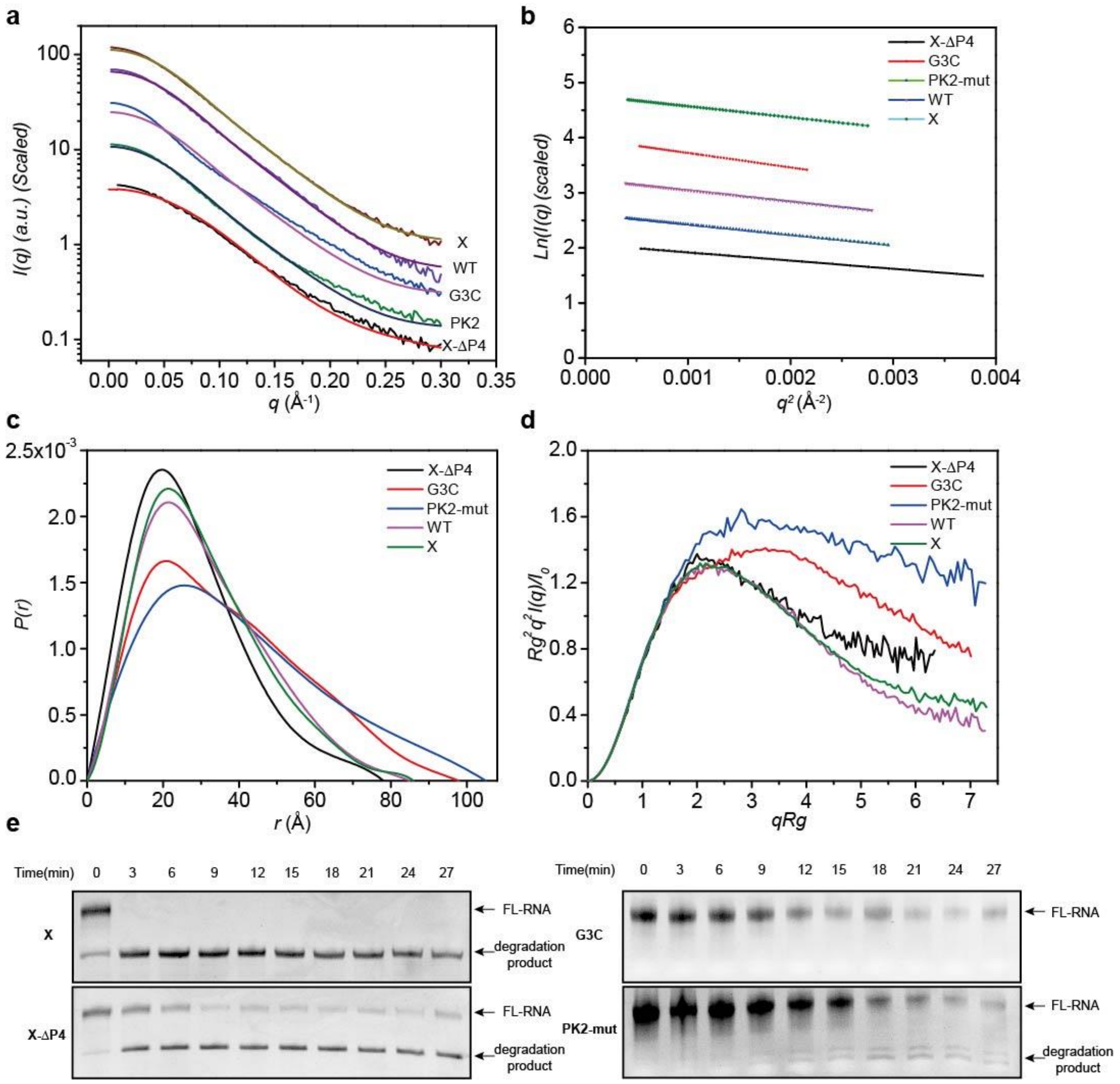

**Figure S10**. Loss of key tertiary interactions disrupts the structural integrity of ZIKV xrRNA1. Experimental scattering profiles (**a**), guinier fitting (**b**), PDDFs (**c**) and dimensionless kratky plots (**d**) for ZIKV xrRNA1 and its various mutants in 5 mM $Mg^{2+}$. In **a**, the theoretical scattering curve of ZIKV xrRNA1 crystal structure was fitted to the experimental scattering curves of xrRNA1, xrRNA1-X, xrRNA1-G3C, xrRNA1-PK2mut, respectively, the theoretical scattering curve of ZIKV xrRNA1 crystal structure without P4 was fitted to xrRNA1-X-ΔP4. (**e**) Xrn1 resistance assay for different ZIKV xrRNA1 mutants. Source data for panel **e** are provided as a Source Data File.

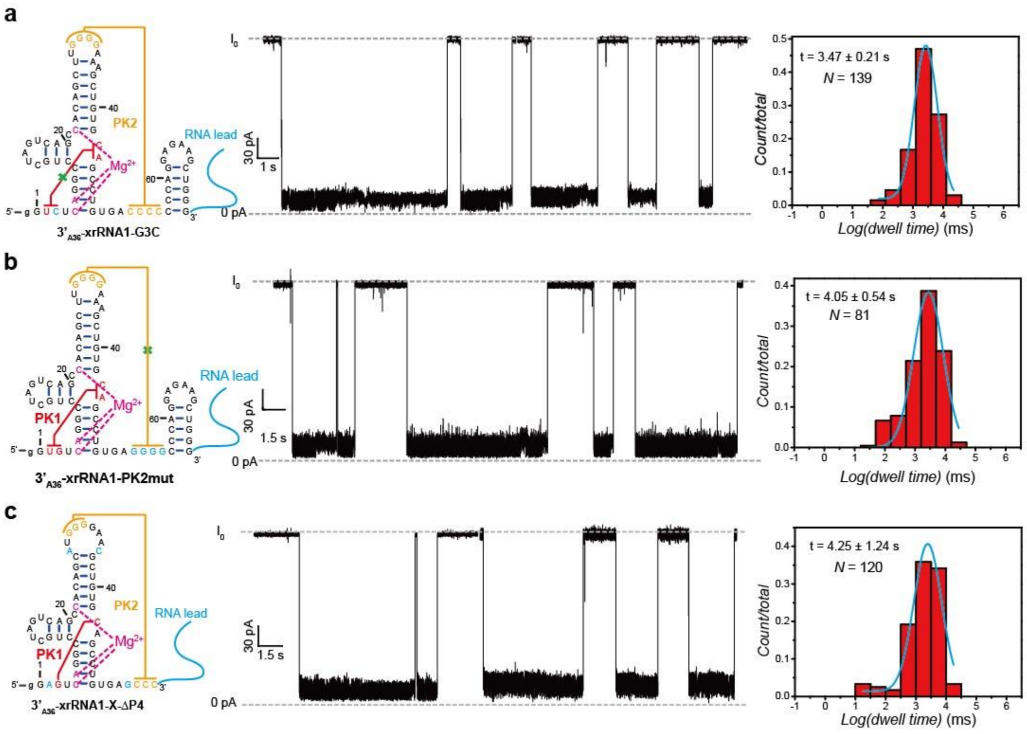

**Figure S11**. 3'→5' directional unfolding of xrRNA1 mutants in the nanopore. (**a-c**) Secondary structure (**left**), representative current blockade traces (**middle**) and dwell time distribution histograms (**right**) for 3'$_{A36}$-xrRNA1-G3C (**a**), 3'$_{A36}$-xrRNA1-PK2mut (**b**) and 3'$_{A36}$-xrRNA1-X-ΔP4 (**c**).

**a**

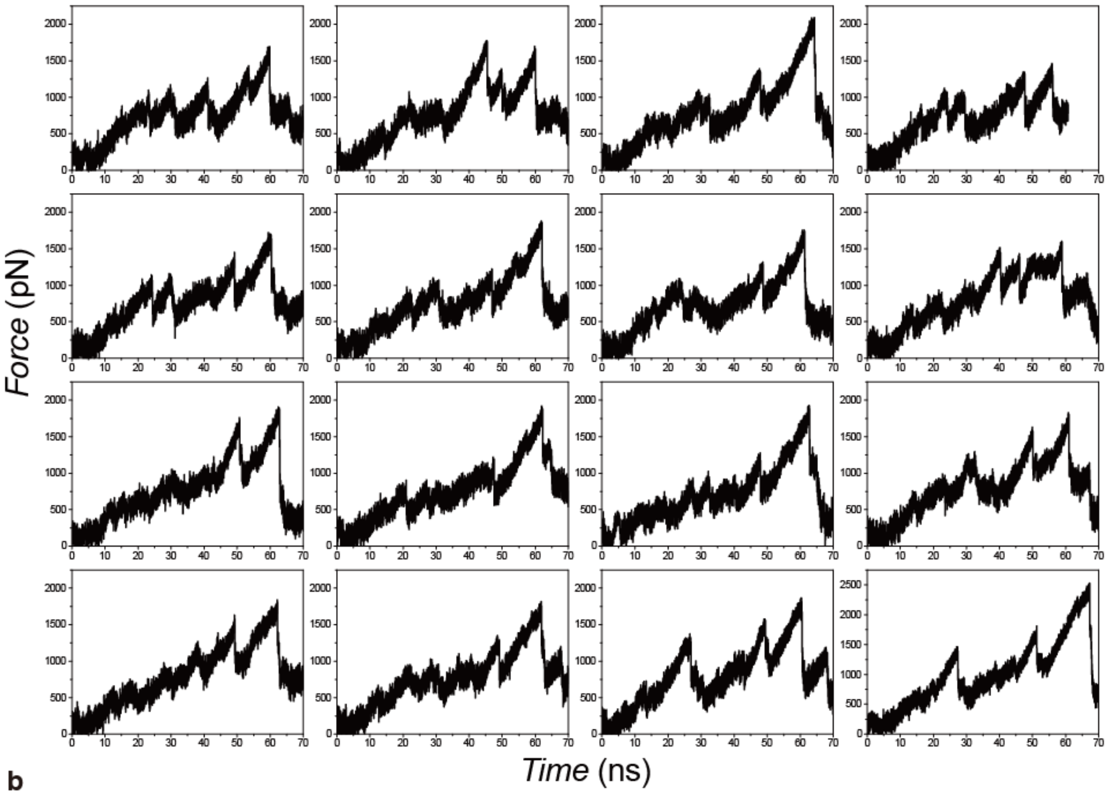

**b**

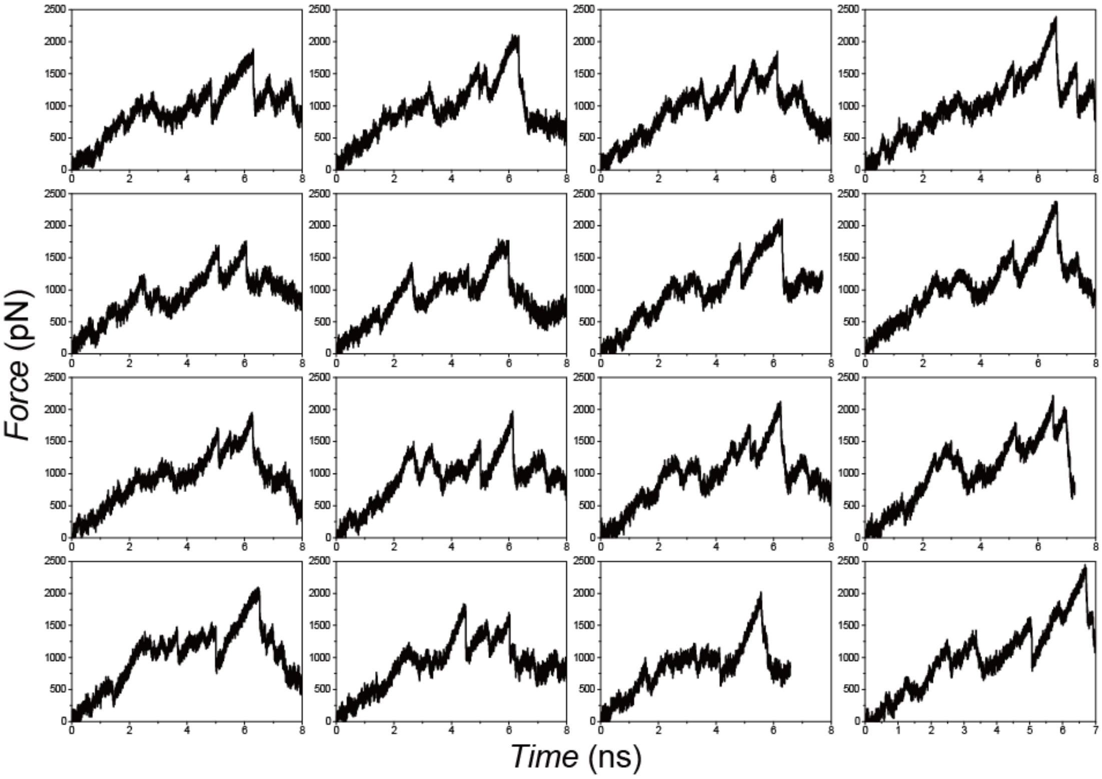

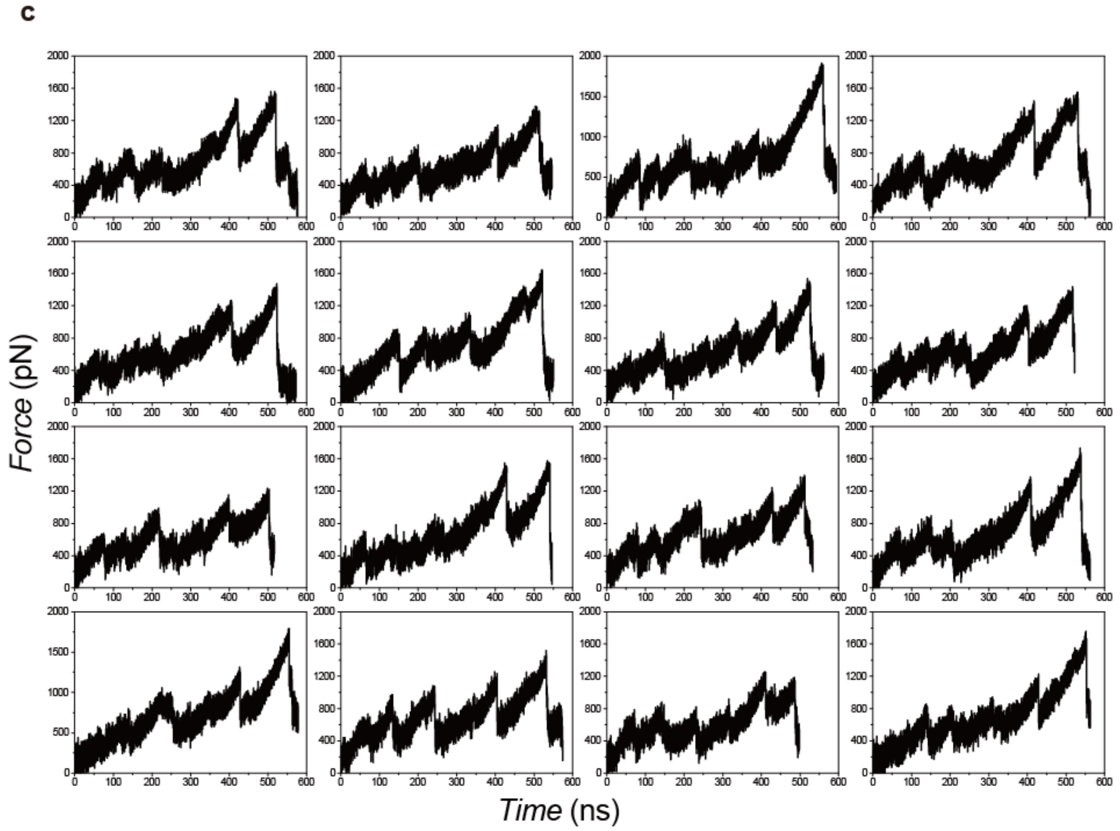

**Figure S12**. Force profiles for 5’-end unfolding of ZIKV-xrRNA1 at pulling rate of 0.1nm/ns (**a**), 1nm/ns (**b**) and 0.01nm/ns (**c**) (sixteen trajectories for each panel). Accordingly, the average height of the largest force peaks are 1713 ± 222 pN, 1940 ± 224 pN and 1387 ± 184 pN; and the average resistance forces over the initial regime (from the entry of 5’-end to total disruption of coordination of magnesium ion with RNA, which is associated to last major force peak) are 839 pN, 1036 pN and 654 pN.

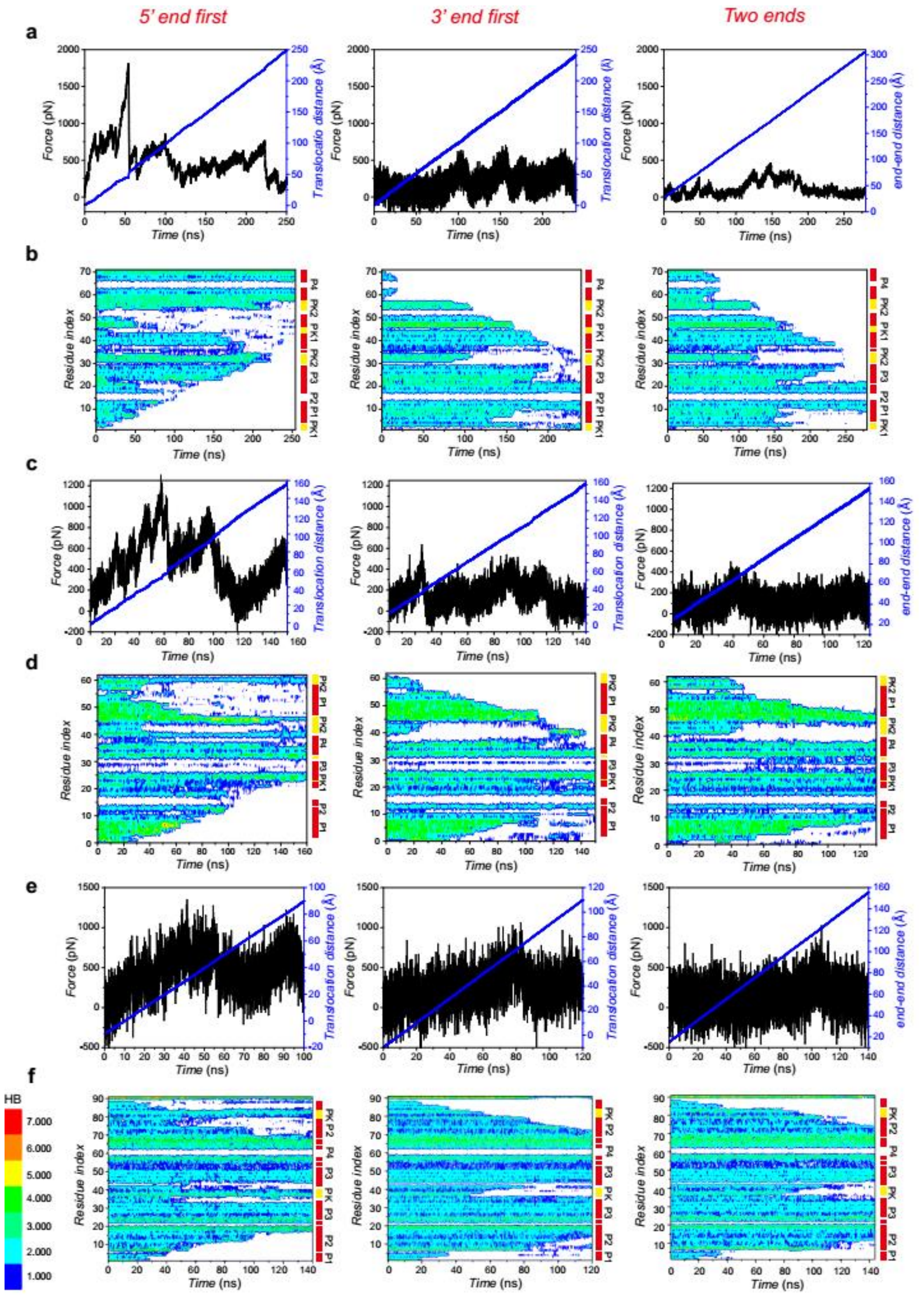

**Figure S13**. Mechanical unfolding of ZIKV-xrRNA1 (**a-b**), HCV-IRES (**c-d**) and THF riboswitch (**e-f**) by MD simulation at pulling rate of 0.1nm/ns. (**a, c, e**) Force profile (black) and pulling coordinate (blue) for RNAs under 5'-end (left) 3'-end (middle) and two-ends (right) pulling directions. (**b, d, f**) Hydrogen bonds (HB) for each residue during mechanical unfolding of three RNAs under different pulling directions. In the case of THF-riboswitch, the hydrogen bonds involved with ligands were also taken into consideration.

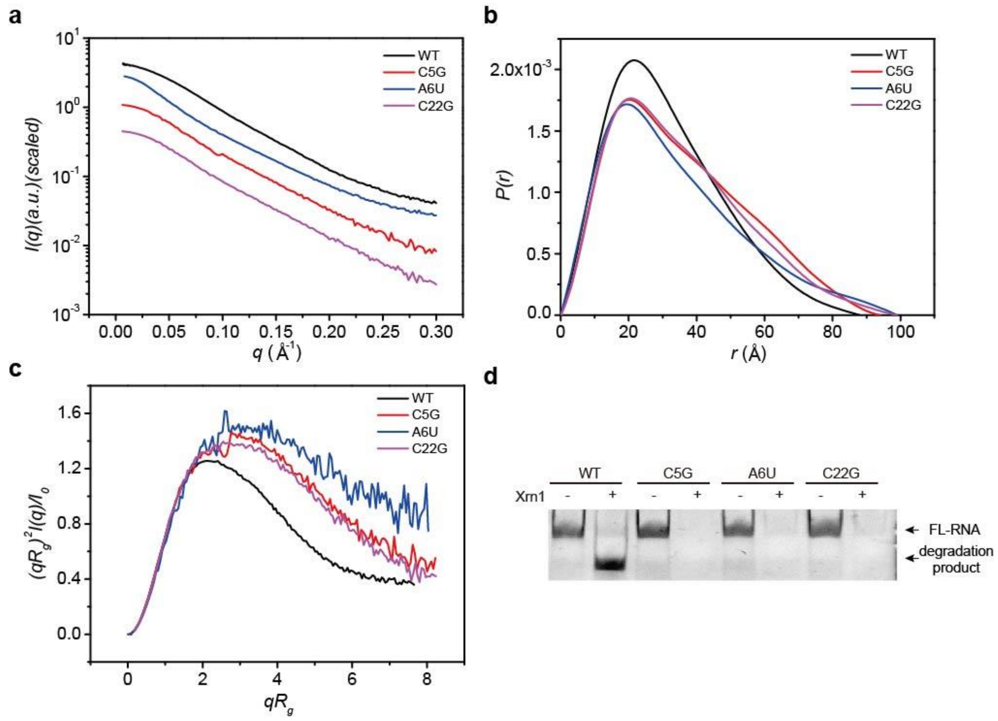

**Supplementary Figure 14**. Mutants that disrupt $Mg^{2+}$ binding site (C5G, A6U and C22G) cause improper folding of ZIKV xrRNA1 and loss of Xrn1 resistance ability. Experimental scattering profiles (**a**), PDDFs (**b**) and dimensionless Kratky plots (**c**) for ZIKV xrRNA1, xrRNA1-C5G, xrRNA1-A6U, and xrRNA1-C22G, respectively. (**d**) Xrn1 resistance assay for ZIKV xrRNA1 and its C5G, A6U, and C22G mutants. Source data for panel **d** are provided as a Source Data File.

**Table S1.** Basic structural parameters for ZIKV-xrRNA1 WT at various $Mg^{2+}$ and $K^+$ concentrations by SAXS.

| Sample | $[Mg^{2+}]$(mM) | ${}^{a}R_g$ (Å) | ${}^{b}R_g$ (Å) | $D_{max}$ (Å) | [c]MW(kD) | [d]MW(kD) |
|---|---|---|---|---|---|---|
| [e]ZIKV-xrRNA1 | 0 | 30.42±1.19 | 32.65±0.33 | 110±2 | 22.44 | 27.62 |
| | 0.001 | 30.39±1.72 | 32.61±0.29 | 110±2 | | 29.61 |
| | 0.005 | 30.31±1.20 | 32.53±0.28 | 108±2 | | 29.56 |
| | 0.01 | 30.26±1.04 | 32.42±0.32 | 108±2 | | 29.38 |
| | 0.02 | 30.02±1.41 | 32.34±0.29 | 107±2 | | 29.38 |
| | 0.05 | 29.92±0.95 | 32.24±0.31 | 107±2 | | 29.61 |
| | 0.1 | 29.64±0.85 | 31.90±0.35 | 106±2 | | 28.78 |
| | 0.2 | 29.22±0.86 | 31.53±0.34 | 105±2 | | 28.65 |
| | 0.4 | 27.18±0.77 | 29.02±0.27 | 98±2 | | 27.51 |
| | 0.6 | 26.38±0.58 | 28.08±0.27 | 97±2 | | 27.13 |
| | 1.0 | 23.34±0.63 | 24.40±0.25 | 86±2 | | 25.84 |
| | 2.0 | 22.53±0.51 | 23.32±0.20 | 80±2 | | 24.93 |
| | 4.0 | 22.38±0.58 | 23.17±0.19 | 79±2 | | 24.79 |
| | 6.0 | 22.49±0.93 | 23.28±0.21 | 81±2 | | 25.07 |
| | 10 | 24.40±0.46 | 25.57±0.37 | 100±2 | | 26.48 |
| [f]ZIKV-xrRNA1 | 0 | 24.45±0.88 | 25.71±0.37 | 95±2 | 22.44 | 22.72 |
| | 0.001 | 24.41±0.84 | 25.64±0.35 | 94±2 | | 22.82 |
| | 0.005 | 24.38±0.77 | 25.58±0.37 | 94±2 | | 23.20 |
| | 0.01 | 24.32±0.64 | 25.60±0.35 | 94±2 | | 22.71 |
| | 0.02 | 24.35±0.76 | 25.45±0.40 | 94±2 | | 22.64 |
| | 0.05 | 24.05±0.51 | 25.38±0.28 | 92±2 | | 22.73 |
| | 0.1 | 23.82±0.65 | 24.95±0.29 | 91±2 | | 22.63 |
| | 0.2 | 23.23±0.62 | 24.27±0.30 | 88±2 | | 22.59 |
| | 0.4 | 22.94±0.52 | 23.95±0.26 | 85±2 | | 22.45 |
| | 0.6 | 22.76±0.48 | 23.74±0.22 | 82±2 | | 22.46 |
| | 1 | 22.66±0.61 | 23.46±0.20 | 80±2 | | 22.39 |
| | 2 | 22.45±0.44 | 23.24±0.22 | 78±2 | | 22.29 |
| | 4 | 22.49±0.61 | 23.29±0.23 | 79±2 | | 22.40 |
| | 6 | 22.67±0.60 | 23.49±0.25 | 82±2 | | 22.48 |
| | 10 | 22.77±0.74 | 23.59±0.25 | 83±2 | | 22.98 |
| [g]ZIKV-xrRNA1 | 0.1 | 23.47±0.30 | 24.16±0.27 | 90±2 | 22.44 | 23.91 |
| | 0.5 | 23.25±0.48 | 24.00±0.27 | 89±2 | | 23.90 |
| | 1.0 | 23.32±0.36 | 24.02±0.25 | 89±2 | | 24.88 |
| | 2.0 | 23.40±0.30 | 24.02±0.23 | 89±2 | | 24.76 |
| | 5.0 | 23.49±0.35 | 24.18±0.26 | 91±2 | | 25.02 |
| | 10 | 24.37±0.27 | 25.30±0.33 | 97±2 | | 25.04 |

[a]derived from Guinier fitting;
[b]derived from GNOM analysis;
[c]MW: molecular weight predicted from sequences;
[d]MW: molecular weight calculated based on the power law of volume of correlation;
[e]Buffer: 20 mM Tris-HCl, pH 7.50;
[f]Buffer: 20 mM Tris-HCl, 100 mM KCl, pH 7.50;
[g]Buffer: 20 mM Tris-HCl, 1000 mM KCl, pH 7.50.

**Table S2.** Transition temperatures for ZIKV-xrRNA1 and its mutants at various buffer conditions by DSC.

| **Sample** | **[$Mg^{2+}$](mM)** | **$^aT_{m1}$(℃)** | **$^bT_{m2}$(℃)** | **$^cT_m$(℃)** |
|---|---|---|---|---|
| ZIKV-xrRNA1 | 0 | 63.33 | 75.30 | - |
| | 1 | 68.07 | 77.63 | - |
| | 2 | 71.41 | 79.29 | - |
| | 3 | 73.92 | 80.86 | - |
| | 5 | - | - | 78.52 |
| | 10 | - | - | 81.25 |
| 5'$_{A36}$-xrRNA1 | 5 | - | - | 77.21 |
| 3'$_{A36}$-xrRNA1 | | - | - | 77.46 |
| xrRNA1-X | | - | - | 75.08 |
| 5'$_{A36}$-xrRNA1-X | | - | - | 73.97 |
| 3'$_{A36}$-xrRNA1-X | | - | - | 73.34 |
| xrRNA1-G3C | | - | 83.22 | - |
| 5'$_{A36}$-xrRNA1-G3C | | - | 81.52 | - |
| 3'$_{A36}$-xrRNA1-G3C | | - | 82.98 | - |
| xrRNA1-PK2-mut | | 77.28 | 83.25 | - |
| 5'$_{A36}$-xrRNA1-PK2mut | | 79.15 | 84.66 | - |
| 3'$_{A36}$-xrRNA1-PK2mut | | 74.98 | 82.60 | - |

[a]Transition temperature derived from the 1st peak, which corresponds to the unfolding of tertiary structures;

[b]Transition temperature derived from the 2nd peak, which corresponds to the unfolding of secondary structures;

[c]Transition temperature corresponds to the cooperative unfolding of xrRNA1.

**Table S3.** Basic structural parameters for ZIKV-xrRNA1 and its mutants at 5 mM $Mg^{2+}$ concentrations by SAXS.

| [a]Sample | [b]$R_g$(Å) | [c]$R_g$(Å) | $D_{max}$ (Å) | [d]MW(kD) | [e]MW (kD) |
|---|---|---|---|---|---|
| xrRNA1-WT | 22.49±0.61 | 23.29±0.23 | 85±2 | 22.44 | 22.40 |
| xrRNA1-X | 24.61±0.14 | 25.50±0.11 | 86±2 | | 25.48 |
| xrRNA1-G3C | 28.05±0.21 | 29.53±0.18 | 98±2 | | 26.57 |
| xrRNA1-PK2mut | 31.92±1.88 | 32.10±0.20 | 105±2 | | 24.22 |
| xrRNA1-X-ΔP4 | 21.11±1.15 | 23.22±2.00 | 78±2 | 17.43 | 17.53 |
| xrRNA1-C5G | 27.94±0.67 | 28.39±0.41 | 94±2 | 22.44 | 26.99 |
| xrRNA1-A6U | 26.97±0.88 | 27.45±1.76 | 99±2 | | 28.42 |
| xrRNA1-C22G | 27.61±0.49 | 27.88±0.43 | 99±2 | | 23.26 |

[a]Buffer: 20 mM Tris-HCl, 100 mM KCl, 5 mM $MgCl_2$, pH 7.50;
[b]derived from Guinier fitting;
[c]derived from GNOM analysis;
[d]MW: molecular weight predicted from sequences;
[e]MW: molecular weight calculated based on the power law of volume of correlation.

**Table S4. The Dwell time of directional translocation through the nanopore for ZIKV xrRNA1 and its mutant constructs.**

| Construct | Buffer condition | Translocation direction | Dwell time (s) |
|---|---|---|---|
| xrRNA1-WT | 5 mM $Mg^{2+}$ | 5' → 3' | > 300 |
| | | 3' → 5' | 2.06 ± 0.58 |
| xrRNA1-X | 5 mM $Mg^{2+}$ | 5' → 3' | 60.78 ± 5.32 |
| | | 3' → 5' | 2.01 ± 0.15 |
| | 5 mM EDTA | 5' → 3' | 0.70 ± 0.02 |
| | | 3' → 5' | 1.21 ± 0.07 |
| xrRNA1-G3C | 5 mM $Mg^{2+}$ | 5' → 3' | 0.09 ± 0.04 |
| | | 3' → 5' | 3.47 ± 0.21 |
| xrRNA1-PK2mut | 5 mM $Mg^{2+}$ | 5' → 3' | 5.49 ± 1.74 |
| | | 3' → 5' | 4.05 ± 0.54 |
| xrRNA1-X-ΔP4 | 5 mM $Mg^{2+}$ | 5' → 3' | 46.77 ± 8.81 |
| | | 3' → 5' | 4.25 ± 1.24 |

**Table S5. The primary sequences of ZIKV xrRNA1 and its mutant constructs used in this study.**

| Construct | RNA sequences |
|---|---|
| 5’$_{A36}$-xrRNA1-WT | 5’-(A)$_{36}$GGUGUCAGGCCUGCUAGUCAGCCACAGCUUGGGGAA AGCUGUGCAGCCUGUGACCCCCCCAGGAGAAGCUGGG-3’ |
| 3’$_{A36}$-xrRNA1-WT | 5’-GGUGUCAGGCCUGCUAGUCAGCCACAGCUUGGGGAAAGC UGUGCAGCCUGUGACCCCCCCAGGAGAAGCUGGG-(A)$_{36}$-3’ |
| 5’$_{A36}$-xrRNA1-X | 5’-(A)$_{36}$GGAGUCAGGCCUGCUAGUCAGCCACAGCAUGGGGAA CGCUGUCAGCCUGUGAGCCCCCCAGGAGAAGCUGGG-3’ |
| 3’$_{A36}$-xrRNA1-X | 5’-GGAGUCAGGCCUGCUAGUCAGCCACAGCAUGGGGAACGC UGUGCAGCCUGUGAGCCCCCCAGGAGAAGCUGGG-(A)$_{36}$-3’ |
| 5’$_{A36}$-xrRNA1-G3C | 5’-(A)$_{36}$-GGUCUCAGGCCUGCUAGUCAGCCACAGCUUGGGGAA AGCUGUGCAGCCUGUGACCCCCCCAGGAGAAGCUGGG-3’ |
| 3’$_{A36}$-xrRNA1-G3C | 5’-GGUCUCAGGCCUGCUAGUCAGCCACAGCUUGGGGAAAGCU GUGCAGCCUGUGACCCCCCCAGGAGAAGCUGGG-(A)$_{36}$-3’ |
| 5’$_{A36}$-xrRNA1-PK2-mut | 5’-(A)$_{36}$GGUGUCAGGCCUGCUAGUCAGCCACAGCUUGGGGA AAGCUGUGCAGCCUGUGAGGGGCCCAGGAGAAGCUGGG-3’ |
| 3’$_{A36}$-xrRNA1-PK2-mut | 5’-GGUGUCAGGCCUGCUAGUCAGCCACAGCUUGGGGAAAGCU GUGCAGCCUGUGAGGGGCCCAGGAGAAGCUGGG-(A)$_{36}$-3’ |
| 5’$_{A36}$-xrRNA1-X-ΔP4 | 5’-(A)$_{36}$-GGAGUCAGGCCUGCUAGUCAGCCACAGCAUGGGGAA CGCUGUGCAGCCUGUGAGCCC-3’ |
| 3’$_{A36}$-xrRNA1-X-ΔP4 | 5’-GGAGUCAGGCCUGCUAGUCAGCCACAGCAUGGGGAACG CUGUGCAGCCUGUGAGCCC-(A)$_{36}$-3’ |

**Table S6. SAXS data collection parameters and software employed for data analysis.**

| ***Data Collection Parameters*** | | |
|---|---|---|
| Facilities and parameters | Settings and values | |
| Beam line | 12ID-B ( APS, ANL) | BL19U2 (SSRF) |
| Wavelength (Å) | 0.8857 | 1.033 |
| Detector | Pilatus 1M | Pilatus 100K |
| $q$ range ($Å^{-1}$) | 0.005-0.89 | 0.009-0.415 |
| Exposure time (s) | 30-60 | 60 |
| Concentration range ( mg/ml) | 0.75-3 | 0.75-3 |
| Temperature (K) | 298 | 298 |
| ***Software Employed*** | | |
| Primary Data Processing | Matlab/BioXTAS RAW/PRIMUS | |
| $P$(r) Function | GNOM | |
| SAXS Profile Computation | CRYSOL | |
| Molecular Visualization | PyMol | |
